# Molecular insights into the pathways underlying naked mole-rat eusociality

**DOI:** 10.1101/209932

**Authors:** Eskeatnaf Mulugeta, Lucile Marion-Poll, David Gentien, Stefanie B. Ganswindt, André Ganswindt, Nigel C. Bennett, Elizabeth H. Blackburn, Chris G. Faulkes, Edith Heard

## Abstract

**Background:** Eusociality is the highest level of social organization and naked mole-rats (NMR)s are amongst the few mammals showing this unique social behavior; nevertheless, little is known about the molecular mechanisms underlying the eusociality of NMRs.

**Results:** Gene expression profiling of NMR brain and gonads (ovary and testis), from animals belonging to different reproductive castes, revealed robust gene expression differences between reproductive and non-reproductive members of NMR colonies. In the brain, dopaminergic pathways appear to be potential players in NMR eusocial behaviour. Breeding animals (queens and breeding males) showed increased expression of genes involved in dopamine metabolism. Using immunohistochemistry, we notably found these differences to be in dopaminergic hypothalamic areas, which provide inhibitory control over the secretion of prolactin, amongst other regions. Furthermore, plasma prolactin concentrations were elevated in many non-breeders (of both sexes), often reaching levels exceeding that of pregnant or lactating queens, suggesting a role for hyperprolactinaemia in socially-induced reproductive suppression. We also found that the ovaries of non-breeding females are arrested at pre-pubertal stage. They contained fewer supporting stromal cells compared to queens, and had very low expression of the aromatase gene *Cyp19A1* (a key enzyme in estrogen synthesis) compared to non-breeding females. In the testes, genes involved in post meiosis spermatogenesis and sperm maturation (*Prm1, Prm2, Odf3* and *Akap4*) were highly expressed in breeding males compared to non-breeders, explaining the low sperm number and impaired sperm motility characteristic of non-breeding males.

**Conclusions:** Our study suggests that extreme reproductive skew, one of the defining features of eusociality, is associated with changes in expression of key components of dopamine pathways, which could lead to hypogonadism and a lifetime of socially-induced sterility for most NMRs.

## Background

Eusociality is the highest level of social organization and is perceived as one of the major transitions in the evolution of life [1, 2]. The original definition, derived from studies of social insects, requires three criteria: a reproductive division of labour, overlapping generations, and cooperative care of young [3, 4], although it has been argued that physically distinct morphological castes should also be present [5]. This extraordinary cooperative way of life is famously observed in invertebrates, such as the social hymenoptera, but also other diverse groups such as crustaceans, and more recently, some mammals (6–9). The importance of eusociality as a strategy is exemplified by the fact that eusocial species may constitute 75% of the insect biomass in some ecosystems [10, 11]. The taxonomic diversity of eusociality has led to much debate about its definition, especially among mammals (for review see [12]), but it is generally accepted that among the African mole-rats (Family: Bathyergidae), the naked mole-rat (*Heterocephalus glaber*) and Damaraland mole-rat (*Fukomys damarensis*) have independently evolved eusociality [8, 13, 14].

NMRs live entirely underground in the arid regions of East Africa in colonies that may contain up to 300 individuals [15]. Most colonies of NMRs have only one breeding female (the queen), who mates with one to three selected males, and the rest of the colony (both sexes) are reproductively suppressed [8, 16]. The reproductive status of the queen and the breeding males may be stable for many years - NMRs can live up to 32 years in captivity [17]. The queen is dominant in the colony social hierarchy, and evidence suggests that she exerts a dominant control mechanism of reproductive suppression over the non-breeders of both sexes, and possibly also the breeding male (18–21). The remaining members of the colony of both sexes are morphologically very similar (externally) and do not exhibit sexual behavior, but perform tasks that are essential for the survival and wellbeing of the colony: foraging, colony defense, maintenance of the tunnel system, and care of the young [8, 16]. It has been estimated that more than 99% of non-breeders never reproduce [16]. However, the extreme socially-induced suppression of reproduction is reversible upon removal of the suppressing cues: if the queen or breeding males die or are removed from the colony, or if non-breeding animals are housed singly or in pairs [18, 20–23].

Non-breeding females remain at pre-pubertal anovulatory state despite attaining adult body size, with small uteri and ovaries, and reduced plasma luteinizing hormone (LH) concentrations that is reflected in lower plasma and urinary progesterone levels [18, 19, 24]. Upon removal of the queen, and often after fighting and competition among non-breeding females, one non-breeding female attains breeding status, and undergoes anatomical, behavioral, and endocrine changes, including elongation of the body, perforation of the vagina, increased dominance and aggression, and activation of the ovaries and ovulation, with resulting increased urinary and plasma progesterone [22, 24–26]. The degree of suppression in non-breeding males is less pronounced than the suppression in non-breeding females in that gametes are produced. Non-breeding males have lower urinary testosterone levels, plasma luteinizing hormone (LH), and lower reproductive tract to body mass ratio compared to breeders [20, 27]. Nevertheless, non-breeding males are able to undergo spermatogenesis and mature spermatozoa are detected in their epididymis and vas deferens, although their number is lower than in breeding males and the majority are non-motile [28]. This indicates that the low concentration of testosterone and LH in non-breeding males is sufficient to support the spermatogenetic cycle, but not the final stages of maturation steps (pituitary FSH may also be inhibited, but to date has not been measured in NMRs). As with females, suppressive effects are reversible upon withdrawal of non-breeding males from their parent colonies and housing singly or pairing with a female - increases in concentrations of urinary testosterone levels and plasma luteinizing hormone (LH) occur rapidly [20].

The detailed mechanism underlying this reproductive suppression remains elusive. The central role of hypothalamic gonadotrophin releasing hormone (GnRH) in integrating environmental cues is well established (29–31), while reproductive development and puberty may be dependent on another hypothalamic peptide, kisspeptin (acting via GnRH) [32]. In NMRs, a lack of priming of the pituitary gland by impaired release of hypothalamic GnRH may be a key component in reproductive suppression [18, 27]. Nevertheless, GnRH is still produced: the number of GnRH-1 immunoreactive cell bodies does not differ between breeding and non-breeding NMRs within or between the sexes [33]. However, breeding females have greater numbers of kisspeptin cell bodies in key areas of the hypothalamus, suggesting that emergence from a socially-induced hypogonadotrophic/prepubertal state in female NMRs may involve kisspeptin-related pathways. Recently, elevated levels of the RF amide related protein 3 (RFRP-3 or GnIH) in the brain of non-breeders have been implicated as a component in the suppression of reproduction in NMRs, through the inhibition of GnRH secretion [34]. It is currently unknown how behavioural and other sensory cues (such as signature odours and vocalisations) mediate the extreme social suppression of reproduction in NMRs.

In recent years, gene expression profiling of several eusocial insects have provided insight into the molecular mechanisms and evolutionary paths to eusociality in insects (35–42). However, although genomic and transcriptomic analyses of NMRs and other mole-rats have also been published, these have focused on individual animals or cross species comparisons (43–45). So far no comprehensive comparative analysis within and among sexes and reproductive castes of NMRs has been reported. Here we use RNA-sequencing (RNA-seq) to undertake the extensive transcriptome profiling of breeding and non-breeding NMR brains and gonads (ovary and testis). This has revealed striking gene expression differences that point to the possible mechanisms underlying eusociality in a mammal, and extreme socially-induced reproductive suppression.

## Results

### Changes in the dopaminergic system are strongly associated with NMR reproductive status

In order to determine the origin of the behavioral and reproductive differences between NMR social classes and elucidate the molecular pathways that contribute to these differences, we performed gene expression profiling using very high coverage RNA-Seq on whole brains of several animals that constitute both sexes and different reproductive classes (*Additional file 1A*: animals used and *Additional file 1B*: RNA-seq information). We first explored the global expression levels, and based on principal component analysis (PCA), the first principal components clearly captured the reproductive caste differences: with the queens and other castes distinctly clustering in their respective groups. Similarly, using a hierarchical clustering technique, non-pregnant NMR queens (Qs) cluster separately to the rest of the colony members (*Supplementary Figure 1A,B*). The rest of the colony members - non-breeding females (NBFs), non-breeding males (NBMs), breeding males (BMs), and a pregnant Q clustered as another group, with the majority of NBFs and NBMs clustering as one group, independent of sex (*Supplementary Figure 1A,B*). In order to define the gene expression differences between brains, we performed differential expression analysis (see Methods) and identified several genes that show significant gene expression differences (FDR<0.05 Benjamini-Hochberg multiple testing correction; a log2 fold change >1; and >1 CPM, counts per million) between breeding and non-breeding animals (*Figure 1A,B*). The number of differentially expressed genes (DEGs) follows social and reproductive status (*Figure 1A,B*), with the highest number of DEGs between breeding animals (Qs vs BMs), or breeding animals compared to non-breeding animals, and with minor differences between subordinate animals (*Figure 1A,B*).

**Figure 1:**
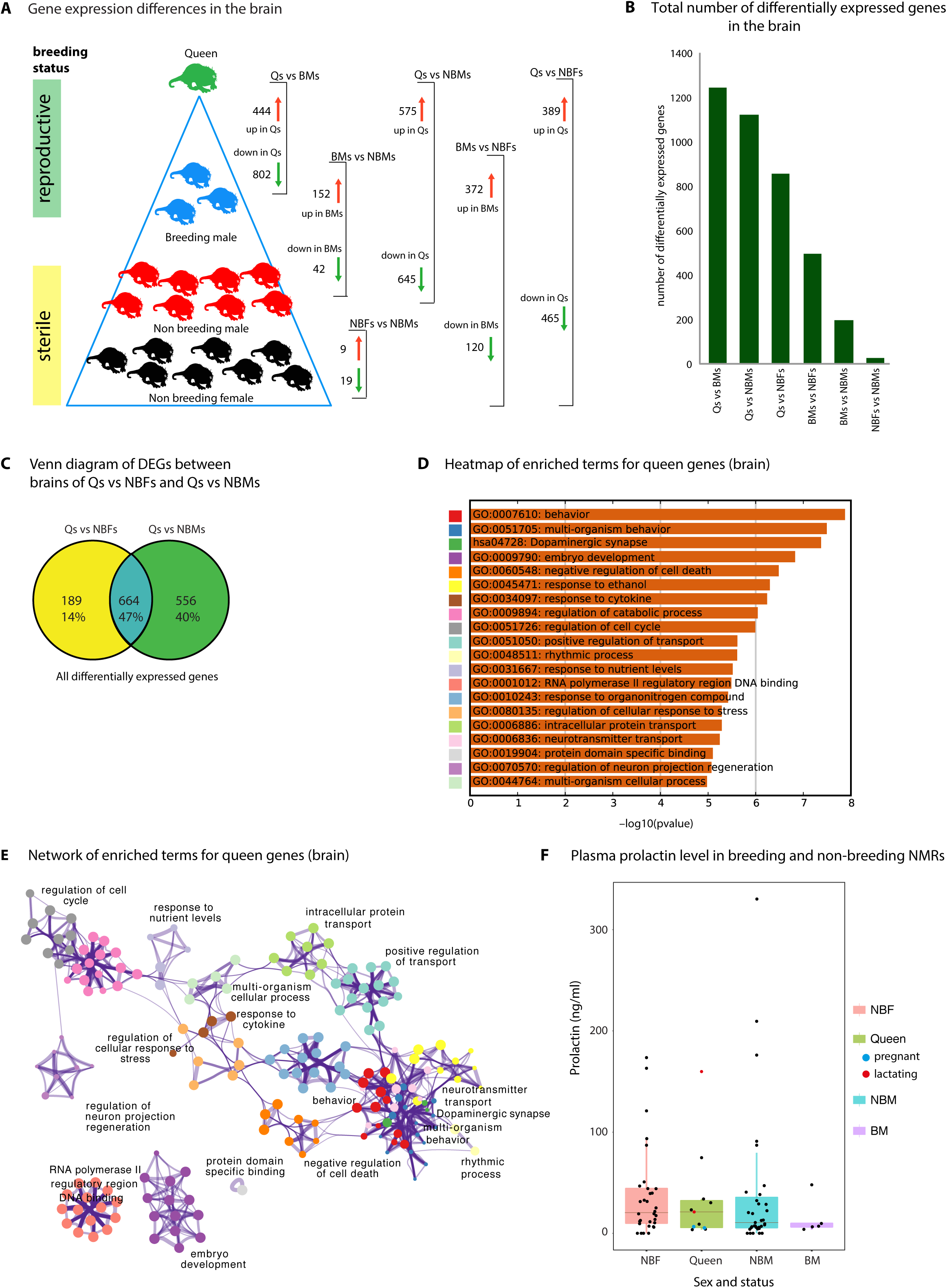
Gene expression profile of naked mole rat colony members. **A)** Number of DEGs between the different members of a naked mole rat colony brains. DEGs (FDR <0.05, log fold change >1) were divided into up regulated (higher expression in one group) and down regulated (lower expression in one group) and the final count is plotted. **B)** Total number of DEGs in each comparison (similar to *Figure 1A*, but total number in each comparison plotted). **C)** Venn diagram showing DEGs that are common in the comparison between Qs vs NBFs and Qs vs NBMs (Q genes, 661 genes in total), and genes that show specific differential expression between Qs vs NBFs (192 genes) and Qs vs NBMs (560 genes). **D)** Heatmap of enriched terms (Canonical Pathways, GO Biological Processes, Hallmark Gene Sets, KEGG Pathway) for genes that show differential expression in the comparison between Qs vs NBFs and Qs vs NBMs (Q genes). Significance of enrichment is indicated on the x-axis in –log_10_(p-value). The color code on the y-axis is used to show the clustering and relation of the significantly enriched networks/terms (shown in *Figure 1E*). Detailed list of enriched terms and families can be found in *Additional file 3.* **E)** Network of enriched terms (*Figure 1D*) colored by cluster ID, indicating the relationship between the different enriched terms (cluster ID color correspond to the color code shown in *Figure 1D* y-axis). Nodes that share the same cluster are close to each other. Each circle node represents a term and the size of the circle is proportional to the number of genes that fall into that term, and the identity of the cluster is indicated by its color (nodes of the same color belong to the same cluster). Similar terms are linked by an edge (the thickness of the edge represents the similarity score). One term from each cluster is selected to have its term description shown as label. **F)** Plasma prolactin concentrations in breeding and non-breeding NMRs in samples taken across thirteen colonies. Abbreviations: Q, queen; Q1_Tr, technical replicate for Q1; NBF, non-breeding female; NBM, non-breeding male; BM, breeding male; DEGs, differentially expressed genes.

To elucidate the nature of these gene expression differences in more detail we examined each group in pairwise comparisons. We first focused on differences between Qs (n=2) and NBFs (n=3). Based on a global gene expression comparison, the Qs and NBFs clearly cluster as two distinct groups, suggesting the clear gene expression difference in the brain of Qs and NBFs (*Supplementary Figure 2A,B*). In total, 854 genes were differentially expressed (389 higher expression in the Qs and, 465 lower expression in Qs) (*Figure 1A,B see Qs vs NBFs, Supplementary Figure 2C, Additional file 2*). These DEGs are enriched for pathway and biological process terms that include synaptic signaling, dopaminergic synapse, positive regulation of transport, and neurotransmitter receptor activity (*Supplementary Figure 2D,E,F,G, and Additional files 3, 4, 5*). Interestingly, genes related to dopamine were particularly enriched in the DEGs. Dopamine is a catecholamine (a monoamine compound derived from the amino acid tyrosine) and has a neurotransmitter function. Several key-genes, such as the tyrosine hydroxylase (*Th*) gene (the rate-limiting enzyme in the synthesis of catecholamines), the vesicular monoamine transporter (*Vmat2* or *Slc18a2*) which transports monoamine neurotransmitters into the vesicles to be released at the synapse, and the dopamine transporter (*Dat* or *Slc6a3*) responsible for the reuptake of dopamine were found to be highly expressed in the Qs compared to the NBFs (*Additional file 2*). *Th* and *Dat* were both in the top 10 of our DEG list, suggesting higher dopamine production in the Qs brain. In addition, the most abundant dopamine receptors *Drd1a*, *Drd2* and *Drd3* were all down-regulated in the Qs, while the *Drd5* receptor gene was up-regulated (*Additional file 2*). Genes related to the other catecholamines adrenalin and noradrenalin (dopamine beta-hydroxylase *Dbh*, phenylethanolamine N-methyltransferase *Pnmt*, and the different adrenergic receptors alpha and beta) were not differentially expressed between Qs and NBFs, indicating that the differences are specific to the dopaminergic system.

Next, we looked at the gene expression differences between BMs and NBMs. As breeding status of male NMRs can be difficult to determine [28], we used two criteria to define their reproductive status in the colony: 1) long-term observational information, and 2) testis to body mass ratio (*Supplementary Figure 3A*). On PCA and hierarchical clustering (using global gene expression), unlike the distinction between Qs and NBFs, BMs and NBMs do not fully resolve into separate clusters (*Supplementary Figure 1B, Supplementary Figure 3B,C*). For animals that fulfill the two criteria above (3 BMs and 3 NBMs, excluding one NBM whose status was uncertain), we performed differential gene expression analysis and identified 193 genes (with 152 showing higher expression in BMs and 42 lower expression in BMs; *Figure 1A B see Qs vs BMs, Supplementary Figure 3 D, Additional file 6*). These DEGs are enriched for pathways and biological processes that are involved in nitrogen compound transport, regulation of nervous system development, synaptic signaling, dopaminergic neuron differentiation, behavior (*Supplementary Figure 3E,F,G,H and Additional files 7, 8, 9*). Similarly to the comparison between Qs vs NBFs, we also found *Th* and *Dat* to be amongst the most highly expressed genes in BMs compared to NBMs (*Additional file 6*), suggesting that similar mechanisms, involving the dopaminergic pathway, may account for the behavioral and reproductive differences observed between BMs and NBMs.

Breeding animals (Qs and BMs) are the most dominant and aggressive in the NMR colony. These animals, Qs and BMs, clearly cluster into independent groups (*Supplementary Figure 1A,B, Supplementary Figure 4A*), and also show the highest number of DEGs (*Figure 1A, B, see Qs vs BMs*). In total 1246 genes show significant expression difference between Qs and BMs (444 higher expression in Qs and 802 lower expression in Qs; *Figure 1A,B, Supplementary Figure 4C, and Additional file 10*). These DEGs are enriched for pathways and biological processes that are involved in protein targeting to membrane, mRNA metabolic process, cellular macromolecule catabolic process, regulation of cellular response to stress and related pathways (*Supplementary Figure 4D,E,F,G,* and *Additional file 11, 12, 13*). The enrichment in these pathways, biological processes, and molecular functions, when breeding females are compared to breeding males, is very different from the findings in non-breeding animals. In contrast to the major gene expression differences in the brains of breeding animals (Qs and BMs), non-breeding animals (NBMs and NBFs) show very similar global expression profiles (*Supplementary Figure 1A,B, Supplementary Figure 5A*), with only very few genes showing significant gene expression differences (only 28 DEGs; *Figure 1A,B see NBFs vs NBMs, Supplementary Figure 5C, Additional file 14*). This is in agreement with previous observations showing the absence of behavioral differences (including sexual characteristics), and sexual differentiation (including similarity in external genitals) between NBMs and NBFs [46].

We went on to identify gene expression profiles that are specific to dominant breeding Qs (Q genes). Q genes were identified by taking genes that show differential expression in the comparison between Qs vs NBFs, Qs vs NBMs (Additional file *15*), and Qs vs BMs (*Figure 1A,B, Supplementary Figure 6 A,B,C*). However, as BMs also have reproductive status and are high in the dominance hierarchy of the colony, we focused on genes that are found in the comparison between Qs vs NBFs and Qs vs NBMs (*Figure 1C, Supplementary Figure 6D,E, Additional file 16*). Around half of the genes that were differentially expressed between Qs vs NBFs are shared with Qs vs NBMs (*Figure 1C*). These Q genes are enriched in pathways and biological process terms that are involved in behavior, synaptic signaling, multi-organism behavior, dopaminergic synapse, neurotransmitter transport, dopamine binding, reproductive behavior and other related terms (*Figure 1D,E, Additional file 17, 18, 19*). We performed a similar analysis to define genes that are unique to the BMs by taking genes that are common in the comparison between BMs vs NBFs (*Additional file 20*) and BMs vs NMBs (*Figure 1A,B, Additional file 21*). In general, while we found fewer such BM genes compared to Q genes (*Supplementary Figure 6F,G,H*), these genes are involved in similar pathways as the Q genes. Similar to the Q genes, BM genes are also enriched for visual perception, nitrogen compound transport, behavior, neuron projection morphogenesis, dopaminergic synapse (*Additional file 22, 23, 24*). In addition, similarly to the Q gene list, *Th* and *Dat* were also present in the BM gene list (unique in BMs vs NBMs and BMs vs NBFs). These results (Q genes and BM genes) highlight the possible role of dopamine pathway in the NMR colony social behavior and reproductive suppression.

Our detailed analysis of NMR brain RNA-seq has identified the dopamine system as a probable key player in the breeding status differences. There are several cell groups in the brain which are potentially dopaminergic (named A8 to A16), linked to different functions [47]. In order to independently confirm the differences in the dopamine system between reproductive castes and to explore their localization, we collected brains from a different batch of animals (*Additional file 1A*). Even though the Qs were bigger in size compared to the other animals (56 ± 16 g versus 34 ± 3 g), all the brains had similar sizes with no obvious neuromorphological differences across the castes. We performed immunostaining against Tyrosine hydroxylase (TH) on several sections along the rostrocaudal axis (*Figure 2A*). The most caudal sections contained the A9 (substantia nigra compacta) and A10 (ventral tegmental area), nuclei that comprise the vast majority of dopaminergic neurons. Their respective main targets, the dorsal striatum and nucleus accumbens, can be seen in the most rostral sections. In addition, we looked at the staining in the hypothalamic nuclei A12, A13 and A14. We observed that across the whole brain, the intensity of the TH staining tended to be higher in breeders than non-breeders, though with inter-individual variability. Strikingly, there was a consistent staining of the hippocampus (and to a lesser extent of the cortex) specifically in all NMR breeders, but absent in non-breeders (*Figure 2B*), indicating catecholaminergic innervation of this region. To our knowledge, such a strong hippocampal staining has never been reported in mice or rats. Importantly, breeders showed significantly more TH expression in periventricular hypothalamic nuclei A12 and A14 (*Figure 2C*), which are precisely the two dopaminergic cell groups known to inhibit prolactin (PRL) secretion, a peptide hormone related to lactation and reproduction [48].

**Figure 2:**
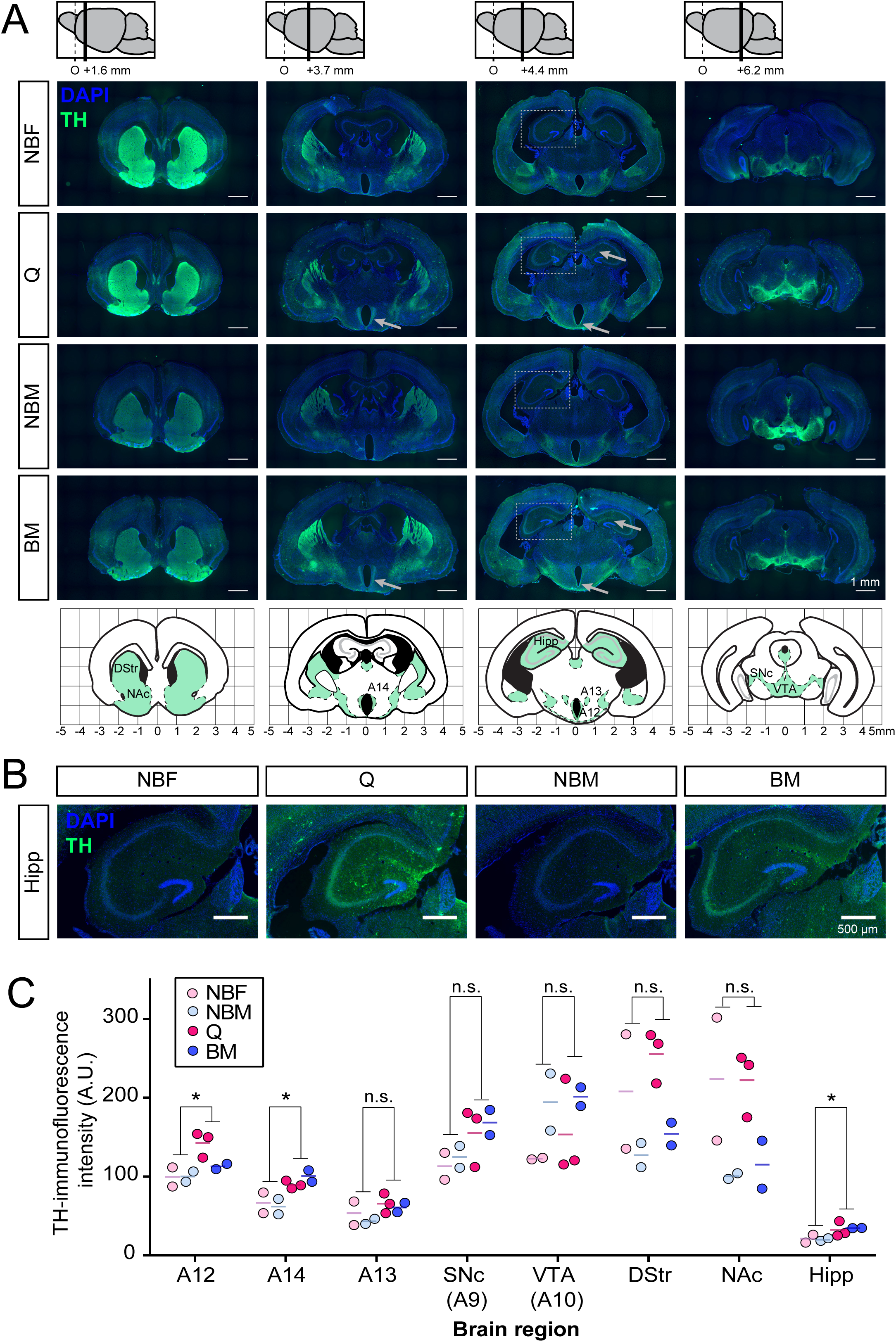
Tyrosine hydroxylase (TH) immunostaining of NMR brain sections. **A)** Representative immunostaining of whole coronal sections for the four reproductive castes (NBF, non-breeding female; NBM, non-breeding male; Q, Queen; BM, breeding male). The top panel indicates the coordinates of the coronal sections relative to O, the origin of slice numbering. Grey arrows point to the regions that show differential staining (quantified in C). Dashed boxes indicate the hippocampal region. The bottom panel represents the regions where TH-staining has been observed. (Labels refer to regions quantified in C: DStr, dorsal striatum; Hipp, hippocampus; NAc, nucleus accumbens; SNc, substantia nigra compacta; VTA, ventral tegmental area). **B)** TH immunofluorescence in the hippocampus. **C)** Intensity of the TH staining in the different brain regions. Non-parametric Mann-Whitney test, *p < 0.05; n.s., non-significant.

To investigate our prediction that elevated prolactin (hyperprolactinemia) may be a component in the suppression of reproduction in non-breeding NMRs, we measured plasma PRL in breeding and non-breeding animals of both sexes in samples taken across thirteen colonies (*Figure 1F*). We found that most samples from non-breeders had detectable concentrations of PRL, and these often reached very high levels: NBF (mean ± SEM) 32.64 ± 6.13 ng/ml; n=44; range, 0.03-173.57 ng/ml; NBM 36.77 ± 9.81 ng/ml; n=49; range, 0.03-330.30 ng/ml. Among breeders, queens had the expected variance in plasma PRL concentrations as part of normal ovarian cyclicity, pregnancy and lactation: 33.02 ± 12.94 ng/ml; n=12; range, 3.60-160.80 ng/ml. Two values were obtained from lactating queens, 21.14 ng/ml (23 days post-partum, at the end of the period of lactation) and 160.80 ng/ml (seven days post-partum), the latter being the highest concentration recorded among the breeding female samples. Although the sample size was low due to the difficulties of identifying them, breeding males had low plasma PRL concentrations, as predicted from our model of suppression: 15.91 ± 6.21 ng/ml; n=7; range, 3.92-47.92 ng/ml.

In conclusion, this comparison of gene expression profiles in NMR brains uncovers differences that follow social status rather than sex, with the Q having a unique gene expression pattern compared to the rest of the colony. Furthermore, we identify dopaminergic pathways as potential key players in NMR eusocial behavior.

### Non-breeding female NMRs have pre-pubertal ovary and lack the production of ovarian estrogen

To characterize the origin of the differential reproductive behavior of Qs and NBFs, we next focused on gene expression differences between their ovaries. Mammalian oogenesis is a highly regulated process that involves several hormonal and molecular pathways. Oogenesis starts in the fetal ovaries with the development of oogonia from primordial germ cells (PGCs). Each oogonium advances until the first stages of meiosis (meiosis I) and become arrested at the prophase stage of meiosis I, forming primary oocytes [49, 50]. After puberty, a few primary oocytes are recruited during each ovarian cycle, and only one oocyte matures to be ovulated. The maturation process involves the completion of meiosis I, and generation of secondary oocytes that are again arrested at the metaphase II stage (meiosis II). The second meiosis will only be finalized after successful fertilization by sperm. Growing oocytes are supported by surrounding somatic cells (follicular cells, granulosa cells, and theca cells) that produce hormones such as estrogens, in response to pituitary gonadotrophic hormones (FSH, LH). At birth, the primary oocytes are embedded into immature primordial follicles, which mature into primary, and secondary follicles. At puberty, oocytes in meiosis II into tertiary and pre-ovulating Graafian follicle [50]. The ovaries of NMR NBFs are reported to be at pre-pubertal stage, with mainly primordial follicles, and may be a few secondary or tertiary follicles [51].

Based on PCA and hierarchical clustering, Qs (n=3) and NBFs (n=3) ovaries clearly group by status and cluster separately (*Figure 3A,B and Supplementary Figure 7A*), supporting previous findings reporting distinct anatomical, endocrine, and physiological differences between the ovaries of Q and NBFs [18, 19, 24]. Irrespective of pregnancy or age, Q ovaries group and cluster together separately from NBFs. Differential expression analysis between the Qs and NBFs (excluding the pregnant Q ovary), specifically identified 1708 genes, with more genes showing significantly lower expression in the Qs compared to NBFs (1175 down regulated genes, lower expression in Qs versus 534 up regulated genes higher expression in Qs compared to NBFs; *Figure 3C, Supplementary Figure 7B, Additional file 25*). DEGs were globally enriched for biological processes and pathways that are related to regulation of nervous system development, response to lipid, meiotic nuclear division, gland development, negative regulation of developmental process, DNA methylation involved in gamete generation and other related terms (*Figure 3D*, *Supplementary Figure 7C,D,E, Additional file 26A, 27A, 28A).* Taking into account the direction of expression changes in Qs versus NBFs, we found that down-regulated genes were enriched for ontology terms related to gamete generation, central nervous system development, developmental process involved in reproduction, meiotic cell cycle process, DNA methylation involved in gamete generation and other related terms (*Supplementary Figure 8A,B,C,D, Additional file 26B, 27B, 28B*). In contrast, up-regulated genes are mainly enriched for inflammatory response, allograft rejection, leukocyte differentiation, positive regulation of cell differentiation, response to lipid, regulation of anatomical structure morphogenesis, regulation of hematopoiesis, cell adhesion and related pathways (*Supplementary Figure 9A,B,C,D, Additional file 26C, 27C, 28C*). These results likely reflect the different cellular composition of Q and NBF ovaries: NBF ovaries are immature and mainly contain primordial and primary follicles (*Supplementary Figure 10*) while Q ovaries develop abundant stromal cells to support oogenesis/folliculogenesis (*Supplementary Figure 10*).

**Figure 3.**
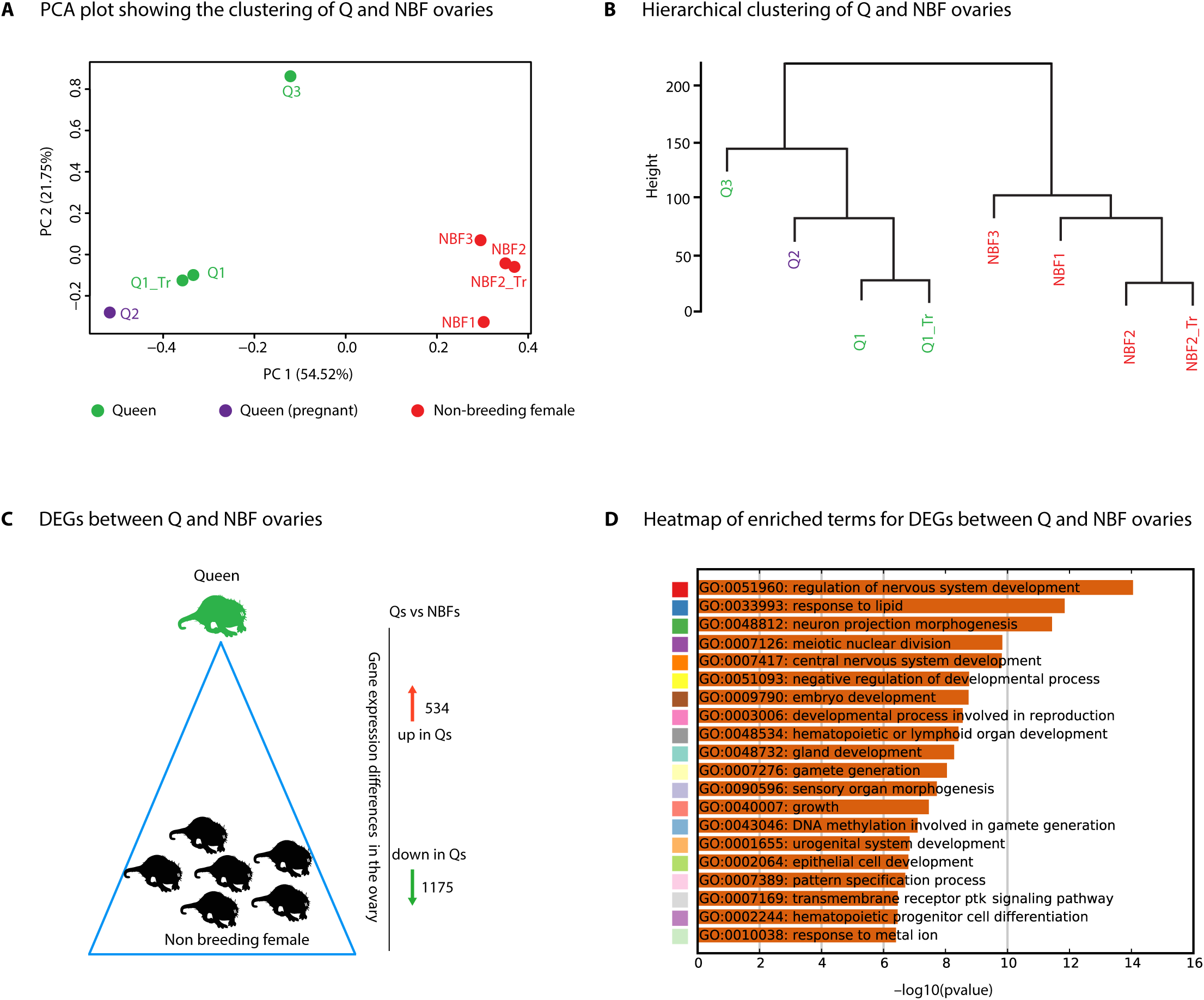
Gene expression profiles of Q and NBF ovaries. **A)** Principal component analysis (PCA) plot showing the clustering of Q and NBF ovaries based on global expression profile. **B)** Hierarchical clustering of ovary samples using euclidean distance matrix computation and ward.D2 agglomeration method. **C)** number of DEGs between Q and NBF ovaries. **D)** Heatmap of enriched terms (Canonical Pathways, GO Biological Processes, Hallmark Gene Sets, KEGG Pathway) for all DEGs colored by p-values. Significance of enrichment is indicated on the x-axis in –log_10_(p-value). The color code on the y-axis is used to show the clustering and relation of this networks (shown in *Supplementary Figure 7C*). More information about the enriched terms can be found in *Additional file 26A*. (Q1_Tr and NBF2_Tr indicate technical replicates for Q1 and NBF2 respectively)

To investigate ovarian differences in more detail, we first compared our data with available transcriptomes of mouse oocytes at different stages of development [52]. We selected two clusters of genes: a first group that show a decline in expression from primary to small antral follicle stages [52] and the other group where expression increased at small and large antral [52] (753 genes in total in the two clusters). However, we did not observe differences in the expression levels of these genes between Q and NBF ovaries (*Supplementary Figure 11A*). This could be due to the fact that, in NMRs, only a few follicles progress to the large antral stage at each reproductive cycle, and thus the difference cannot be quantified from whole ovary expression data. As NBF ovaries have been reported to be in an immature state [18, 19, 24], we next integrated available transcriptomes of whole ovaries from pre-puberal and adult mice [53]. Surprisingly, Q and NBF ovaries did not significantly differ in the expression levels of genes that are differentially expressed between pre-pubertal and adult mouse ovaries (*Supplementary Figure 11B*). However, by focusing on a selection of oocyte-specific genes (*Zp1, Zp2, Zp3, Figla, Nobox, Sohlh1, Gja4, Pou5f1, Oosp1, H1foo, Lhx8, Sohlh2, Kit, Kitl, Nlrp14, Bmp15*), we observed a consistent increase in NBF compared to Q ovaries (*Supplementary Figure 11C*). This is in agreement with our histological assessment (*Supplementary Figure 10*): Q ovaries have more stromal, somatic cells than NBF ovaries, which are enriched in primary oocytes. Upon further examination of the NMR ovary RNA-seq data, in agreement with the enrichment of hormone-producing stromal cells in Q ovaries, we found that the main difference between ovaries of the Qs and NBFs was in the ability to synthesize estrogen. More specifically, among the top-10 list of DEGs, we found that *Cyp19A1 (Aromatase)* - a key enzyme in the synthesis of estrogen- was highly over-expressed in Q compared to NBF ovaries (4-fold change in log2 scale with almost no expression in NBF; *Supplementary Figure 11D)*. Aromatase is important for transformation of androstenedione to estrone and testosterone to estradiol [54, 55]. In addition to *Cyp19A1,* other genes that are involved in steroid hormone biosynthesis (*Cyp7a1, Akr1c18, Akr1d1, Ugt1a1*) were also significantly more expressed in Q ovaries.

As a whole, our RNA-seq analysis suggests that Q ovaries specifically express genes that are required for estrogen production, and thereby undergoes oogenesis/folliculogenesis, ovulation and other sexual differentiation processes. In contrast, NBFs seem to have functioning ovaries that are arrested at a prepubertal stage, and do not have the ability to produce estrogen, explaining their failure to ovulate and subsequent reproductive incompetency.

### Non-breeding male NMRs show signs of defective post meiosis spermatogenesis and sperm maturation

We next wanted to investigate the nature of the gonadal differences between breeding versus non-breeding male NMRs. Mammalian spermatogenesis begins post-natally at puberty, and continues throughout life. A pool of spermatogonial stem cells undergo mitotic divisions to produce spermatocytes, that will undergo meiotic divisions to produce haploid spermatids, which will then undergo further maturation to become mature spermatozoa (56–58). This process of spermatogenesis occurs in seminiferous tubules supported by Sertoli cells and Leydig cells, and coordinated by endocrine signals. Sertoli cells nourish and provide structural support for the developing sperm cells, upon activation by follicle-stimulating hormone (FSH) that is secreted by the pituitary gland, which is in turn under the control of the GnRH secretion from the hypothalamus [58, 59]. Leydig cells that are located outside the seminiferous tubules produce and secrete testosterone and other androgens that are important for spermatogenesis, secondary sexual characteristics, development, and testis volume. Leydig cell function is modulated by the pituitary gonadotrophin luteinizing hormone (LH) [58, 59].

All male NMRs, independently of their breeding status, have been reported to undergo spermatogenesis [19, 26, 27]; however, sperm numbers and motility vary between BMs and NBMs [26, 27]. In order to investigate the molecular pathways that contribute to these differences in spermatogenesis between BMs and NBMs, we analyzed the gene expression profile of whole testes from BMs and NBMs, using the two criteria described above in the brain analysis section to define their status (long-term observational information and testis to body mass ratio). Based on PCA and hierarchal clustering, unlike the clear separation observed for the ovaries of Qs and NBFs, NBM testes do not show a distinct clustering and grouping based on breeding status, with BM3 clustering with non-breeding animals (*Figure 4A,B, Supplementary Figure 12A*). Nevertheless, differential expression analysis identified 780 genes that showed significant changes between BMs and NBMs (522 up-regulated and 258 down-regulated genes in BMs compared to NBMs; *Figure 4C, Supplementary Figure 12B, Additional file 29*). Enrichment analysis for biological processes, molecular functions, and pathways did not highlight obvious enrichment of spermatogenesis genes, but rather for terms such as: regulation of anatomical structure morphogenesis, response to steroid hormone, and embryo development (*Figure 4F, Supplementary Figure 12C,D,E Additional file 30A, 31A, 32A*). By focusing on DEGs that show lower expression in breeding compared to non-breeding testes, we found enrichment for terms that are related to positive regulation of cell-substrate adhesion, positive regulation of cell-substrate adhesion, processes regulation of histone acetylation (*Supplementary Figure 13A,B,C,D, Additional file 30B, 31B, 32B*). However, interestingly, for DEGs that are up-regulated in the testes of BMs we found enrichment of terms related to regulation of anatomical structure morphogenesis, response to steroid hormone, and multicellular organism reproduction (*Supplementary Figure 14A,B,C,D, Additional file 30C, 31C, 32C).* In particular, this last ontology category (GO:0032504) contains a number of genes (45 genes, *Additional file 30D*) involved in key biological processes related to male reproduction, such as meiotic cell cycle, male genitalia development, gamete generation, spermatogenesis, and sperm motility (*Lhcgr, Prm1*, *Prm2*, *Akap4*, *Odf3,* Akap4, *Wfdc2*, *Fdc2*, *Txnrd2, Txnrd3*, *Spata6*, *Spata22*, *Rara*, *Nr2c2*, *Hsf2bp, Cylc2, Cylc1*, *Celef3*, *Ccin*, *Alkbh5*, *Adam29, Atp2b4*; *Additional files 30D*). From this list of 45 genes, *Lhcgr* encodes the receptor for both LH and choriogonadotropin [60, 61]. Binding of LH to its receptor stimulates testosterone production by Leydig cells to promote extra-gonadal differentiation and maturation [62, 63]. *Prm1* and *Prm2* genes encode Protamine proteins that replace histones in later stages of spermatogenesis (sperm elongation) to allow denser packing of DNA (64–66). In mouse models, abnormal expression of *Prm1* and *Prm2* causes a decrease in spermatozoa number, abnormal spermatozoa morphology and motility, damaged spermatozoa chromatin, and infertility (67–73). *Odf3* is transcribed more specifically in spermatids, and is suggested to provide the elastic structure of sperm protecting from damage during epididymal transport [74, 75]. *Akap4* is transcribed only in the post-meiotic phase of spermatogenesis and is a cytoskeletal structure present in the principal piece of the sperm flagellum [76]. Targeted disruption of *Akap4* results in abnormal flagella, and hence motility of spermatozoa [76, 77].

**Figure 4:**
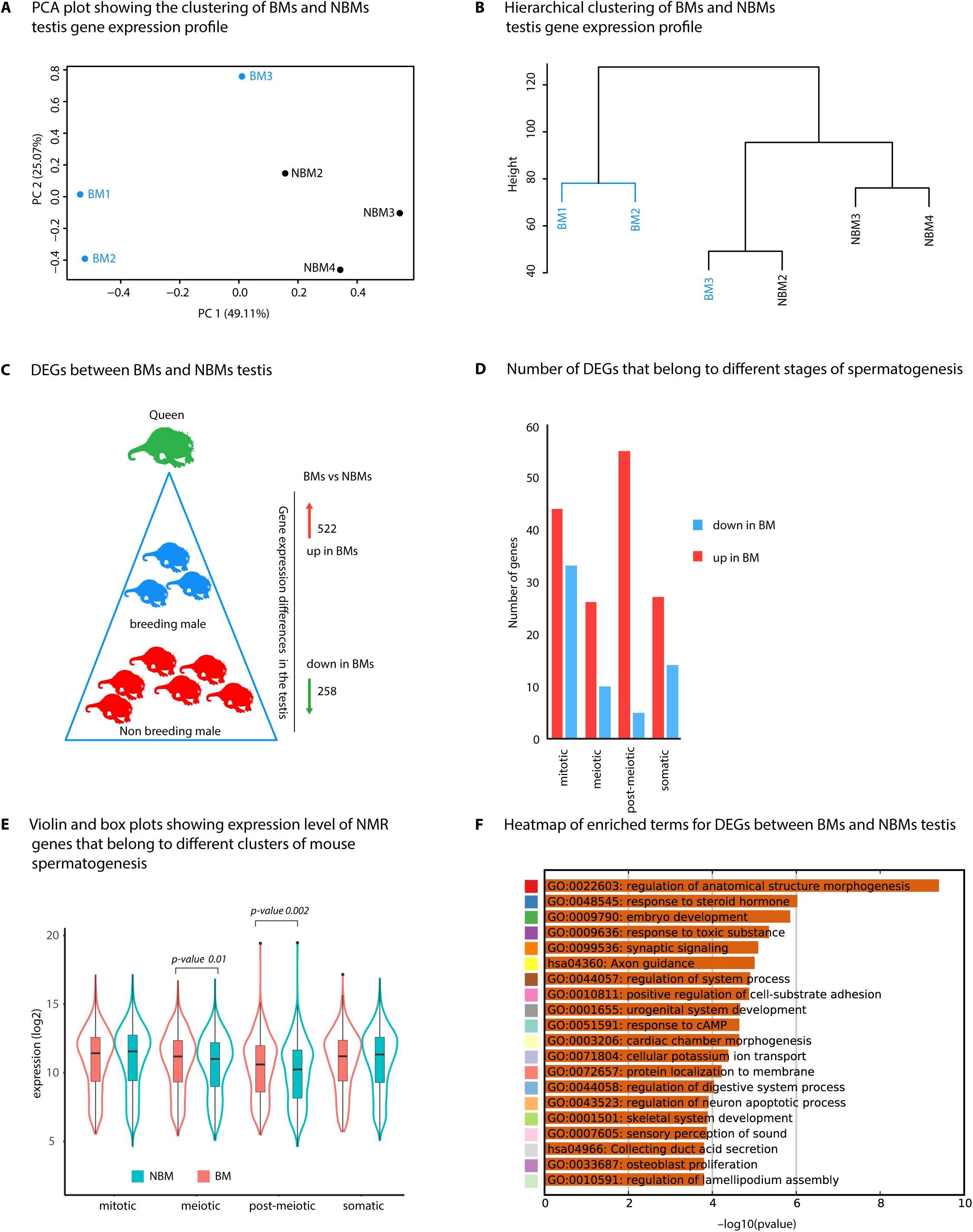
Gene expression profile of breeding and non-breeding males testes. **A)** Principal component analysis (PCA) plot showing the clustering of BMs and NBMs testis based on global gene expression profile. **B)** Hierarchical clustering of testis samples using euclidean distance matrix computation and ward.D2 agglomeration method. **C)** Number of DEGs between breeding and non-breeding animals. **D)** Bar plot showing the number of NMR DEGs (BMs vs NBMs testes) that belong to different stages of mouse spermatogenesis. Gene clusters that are expressed at different stages of mouse spermatogenesis were obtained from Germonline, and mapped to the DEGs that were identified in the comparison between breeding and non-breeding NMR testes. The DEGs which map to different stages of spermatogenesis were divided into up or down regulated, (if they show higher or lower expression in BM respectively) and plotted. **E)** Violin and box plots showing the average expression level of all BMs and NBMs genes (NMR testes) that were mapped to genes which belong to the different clusters (4 clusters) of mouse spermatogenesis. Gene clusters that are expressed at different stages of mouse spermatogenesis were obtained from Germonline, mapped to NMR BMs and NBMs testis expression data, and plotted (x-axis, cluster name; y-axis: average expression level in log_2_ scale). **F)** Heatmap of enriched terms (Canonical Pathways, GO Biological Processes, Hallmark Gene Sets, KEGG Pathway) for all DEGs colored by p-values (generated using Metascape). Significance of enrichment is indicated on the x-axis in –log_10_(p-value). The color code on the y-axis is used to show the clustering and relation of this network (shown in *Supplementary Figure 12C*). Detailed list of enriched terms and genes that belong to these terms is provided *in Additional file 30A*.

To further document what stage of spermatogenesis might be interrupted in NBMs, we imported the list of spermatogenesis-related gene clusters that show specific expression at several different stages of mouse spermatogenesis (mitotic, meiotic, post-meiotic, and somatic clusters,) from the Germonline database (*Additional file 30*) [78]. DEGs belonging to all four clusters tended to show higher expression in BMs than in NBMs, but this difference was more pronounced for post-meiotic genes (*Figure 4D*). More specifically, 10% (55 out of the 522) of up-regulated DEGs in BM belonged to the post-meiotic cluster, while only 2% of down-regulated DEGs in BM (5 out of 258) belonged to this category. This suggests that the main difference in BM and NBM spermatogenesis is related to genes that are important in post-meiotic stages. In addition, we plotted the expression level of all genes that belong to these four clusters. Genes that belong to both the meiotic and post-meiotic stages of spermatogenesis showed significant up-regulation in BMs compared to NBMs (p-value 0.01 and 0.002 respectively, Wilcoxon rank sum test; *Figure 4E*), but this trend was more pronounced for the post-meiotic genes (*Figure 4E*). Finally, to understand the actual nature of the block in spermatogenesis in NBMs, we examined histological sections of BM and NBM testes. Confirming previous observations [19, 26, 27], all stages of spermatogenesis could be observed in both BM and NBM animals (*Supplementary Figure 15A, B*), although it was not possible to obtain a precise quantitative information by this method. Interestingly, a very clear difference was observed in interstitial (Leydig) cell content, with much higher numbers in the BM compared to NBM testes (*Supplementary Figure 15C,D*).

In conclusion, our detailed analysis of NMR testes revealed that although both breeding and non-breeding males undergo spermatogenesis, non-breeders show signs of impaired post-meiotic sperm maturation at the transcriptomic level. This may lead to lower sperm numbers and impaired motility that could underlie their incapacity to breed.

## Discussion

In this study, we set out to define the molecular pathways that might underlie one of the principal features of eusociality, namely the extreme socially-induced suppression of reproduction (reproductive skew) in NMRs. We achieved this by comparison of brain and gonad transcriptomes in the different individuals that make up the NMR social hierarchy, namely Qs (the only breeding females), NBFs, BMs and NBMs. The comparative analysis of gene expression in the brain of different NMR social classes revealed that: 1) the Q has a distinctive gene expression profile when compared to the rest of the colony; 2) both sexes of non-breeding animals have nearly identical gene expression profiles in the brain; 3) several genes show significant gene expression differences in comparisons between breeding animals (Qs vs BMs) and between breeding animals versus non-breeding animals; 4) the Qs and NBFs have clear gene expression differences in the brain, but this is not reflected in comparisons between BMs and NBMs; 5) the dopaminergic pathway is identified as a major pathway that may be linked to the differences between social classes of NMR colonies. More specifically, differences in TH expression (the rate limiting enzyme for the synthesis of dopamine) were found in the hypothalamic areas of the brain which control prolactin secretion [48]. Also, the catecholaminergic innervation (potentially dopaminergic) in the hippocampus was strikingly high in breeders and showed a clear difference between breeders and non-breeders. In mice or rats, there are sparse catecholaminergic afferents, which arise mainly from the ventral tegmental area [79] and the locus coeruleus [80]. In NMR breeders, the origin of these afferents and their role in the different behaviors between castes remains to be investigated. We believe that this potential role of dopamine in NMR eusociality is a significant discovery that is consistent with findings in other eusocial animals. Indeed, the dopaminergic pathway is a highly conserved system across vertebrates and some invertebrates, with many important functions in the nervous system (81–86). These include a crucial role in the control of movement and reinforcement learning. Dopamine is also implicated in sexual arousal, aggression, dominance, sleep, attention, working memory, hormonal regulation, and other functions [82, 87–90]. In eusocial insects and termites, the importance of dopamine in aggression, dominance and social hierarchy has been documented. Brain dopamine levels are elevated in dominant individuals of *Polistes* paper wasps (*Polistes chinensis*), the worker-totipotent ant (*Harpegnathos saltator)*, and the queenless ant (*Diacamma sp.* from Japan); and a positive correlation was observed between ovarian activity and the level of dopamine in wasps (*Polistes chinensis*), bumble bees (*Bombus terrestris*), honeybees (*Apis mellifera L.*), ants (*Harpegnathos saltator*) (91–96). In the honeybee (*Apis mellifera*), tyrosine or royal jelly-fed workers had higher brain dopamine levels than sucrose fed individuals, resulting in ovarian development and the inhibition of foraging in the former [97]. Dietary and topical applications of dopamine in queenless ant subordinate workers have also been shown to induce oocyte growth/activation [93, 96]. In honey bee (*Apis mellifera*), queen pheromone have been shown to modulate dopamine signaling pathways in the worker bees [98]. Furthermore, in male honeybees, juvenile hormone (JH), which regulate development and reproduction, has been shown to increase the levels of dopamine in the brain [99, 100]. JH and brain dopamine levels have been shown to increase during sexual maturation, and topical application of a JH analog increases the level of dopamine in the brain of male honeybees [101, 102]. These investigations indicate that the dopaminergic pathway can be crucially involved in the social system, behavior, and reproductive suppression of many eusocial insects and termites. Our findings point to the exciting possibility that dopamine-mediated mechanisms are also involved in maintaining eusociality in the NMRs, suggesting that these pathways may represent a rather universal strategy for eusociality across the Animal Kingdom.

The potential functional significance of the observed differences in dopamine pathways between the Q and the non-breeders could be at several levels. One possible and well-established mechanism is the action of dopamine on prolactin regulation (reviewed in [48, 103]). In this pathway, dopamine (also known as prolactin inhibiting factor), modulates the release of prolactin from the pituitary gland, and prolactin itself acts on the hypothalamus to regulate the release of GnRH (*Supplementary Figure 16B,C*) [104]. The plasma prolactin data, measured for the first time in this species, fits our model of elevated PRL suppressing reproduction in non-breeders, because plasma PRL concentrations measured in spot samples collected from most non-breeding NMRs often greatly exceeded values which would be considered clinical hyperprolactinemia in humans (*Figure 1F*). For example, levels of circulating PRL between 3 and 15 mg/l are considered necessary for maintaining normal reproductive function, and levels below and above are associated with an increased rate of infertility[105] - generally this equates to fasting concentrations above 25 ng/ml for women and 20 ng/ml for men [106]. Furthermore, plasma PRL in non-breeders often also exceeded that of lactating queens (*Figure 1F*). We did fail to detect PRL in some plasma samples from non-breeders. However, plasma PRL concentrations are known to exhibit circadian variations, and any such pattern or synchrony amongst NMRs remains to be investigated in detail to avoid sampling at times of low secretion. In mammals, hyperprolactinaemia is well known to be a major cause of infertility, and plays a role in this context in lactational suppression of reproduction [107]. A higher level of dopamine in the Q or BM is expected to decrease the release of prolactin. In turn, the absence or decreased levels of prolactin may result in the normal pulsatile secretion of GnRH from the hypothalamus, and subsequent release of LH and FSH from the pituitary gland. LH and FSH exert their effects at the level of the gonad, stimulating sexual differentiation, follicular development, spermatogenesis and release of the sex hormones estrogen (in females) and testosterone (in males) (*Supplementary Figure 16B)*. The low dopamine in non-breeding animals on the other hand, results in elevated levels of prolactin, inhibiting the release of GnRH, and thus LH, FSH, estrogen, and testosterone, and follicular development and spermatogenesis (*Supplementary Figure 16C)*. The inhibitory effect of dopamine on prolactin has been shown in many animals, and specific dopamine neurons (A12 and A14 cell groups, *Supplementary Figure 16A*) are known to regulate the release of prolactin. Interestingly, the unique presence of cell bodies expressing RFRP-3 in the arcuate nucleus of NMRs, reported by [34] may be of significance, as it could potentially allow interaction with the A12 dopaminergic cell groups that regulate PRL section. Despite the differences in expression of genes involved in increasing dopamine in the brain, we did not observe differences in the GnRH expression between breeding and non-breeding NMR brains (both males and females). However, this would fit with predictions based on previous NMR studies that showed similar numbers of immunoreactive GnRH-1 cell bodies among breeders, non-breeders, males and females [33]. This implies that GnRH is produced by all status groups, but only actually released in breeders, due to a block to its secretion in non-breeders. The neuropeptide kisspeptin may play a role in this process among females. Kisspeptin is well known to influence the hypothalamo– pituitary–gonadal axis by direct actions on GnRH-1 neurons [33]. Zhou *et al.* found that breeding females NMRs had increased numbers of kisspeptin immunoreactive cell bodies in the anterior periventricular nucleus and rostral periventricular region of the third ventricle (RP3V). This suggests a role for kisspeptin in the hypogonadotrophic state in female NMRs, acting via mechanisms similar to those that underlie puberty and seasonal breeding in other species. Our observation of small increases in expression of *Kiss-1* and its receptor (*Kiss-1R*) (the latter being statistically significant) in Qs versus NBFs supports this hypothesis. Furthermore, elevated prolactin is known to have a suppressing effect on kisspeptin (and hence GnRH), which would fit a model of suppression involving dopamine pathways. Brown *et al.* (2014) showed that administration of prolactin to mice caused *Kiss-1* mRNA to be suppressed in the RP3V (the region with increased kisspeptin immunoreactive cell bodies in Q NMRs). It remains to be determined how the elevated levels of RFRP-3 (GnIH) in the brain of non-breeders, reported by [34], act in this mechanism that ultimately inhibits GnRH secretion. Interestingly, elevated prolactin has been implicated in many studies as a factor mediating both parental and alloparental care, affiliative and other sociosexual behaviours, in birds and mammals, including rodents and primates [108]. It is thus tempting to speculate that elevated prolactin in NMRs may play a central role in both cooperative behaviour and reproductive suppression.

The hypothalamic block to reproduction in NMRs ultimately results in the well-documented lack of gonadal development in both male and female non-breeders [18, 19, 27]. Not surprisingly, given the large anatomical differences, gene expression profiling of the ovary identified several DEGs between the Qs and NBFs. One of the important differences in gene expression was the significantly higher expression of aromatase in Qs ovary (top10 DEGs), the key enzyme in the production of estrogen. This difference in aromatase expression indicates the ability of the Qs ovary to produce estrogen, and the low production of ovarian estrogen in NBFs. Aromatase knockout mice display underdeveloped external genitalia and uteri, and precocious depletion of ovarian follicles, and anovulation, with development of the mammary glands approximately that of prepubertal WT female mice (109–111). These features of aromatase knockout mice mimic most of the anatomical and physiological and endocrine differences that were observed between the Qs and NBFs of NMR [18, 19, 24]. The enrichment analysis on DEGs also highlights the difference between the Q and NBF ovaries. The small, pre-pubertal ovaries of NBFs mostly contain primordial and primary follicles that are arrested at the meiosis I prophase. In contrast, the fully functional ovary of the Qs contains follicles at all stages of development, including some polyovular follicles that may contain up to three oocytes [112], and larger numbers of stromal cells (*Supplementary Figure 10*). The ovarian stroma is a diverse mix of cell types that includes theca–interstitial cells, immune cells, blood vessels, smooth muscle cells, and several types of extracellular matrix proteins (113–118). The increased stromal cell content in Qs is reflected in the enrichment of GO terms (such as regulation of leukocyte differentiation, regulation of hematopoiesis, cell adhesion) for DEGs unregulated in the Qs. The NBF ovaries, with mostly primary follicles containing oocytes arrested at meiosis I, is reflected in GO terms (such as DNA methylation involved in gamete generation, sexual reproduction, meiotic nuclear division, meiotic cell cycle) for DEGs upregulated in the NBFs. Taken together, the expression analyses reported here is consistent with the observation that the Qs ovary is able to undergo oogenesis/folliculogenesis, ovulation and other sexual differentiation processes; whereas NBF ovaries are arrested at a pre-pubertal stage, lack the ability to produce estrogen, ovulate and undergo sexual differentiation (*Supplementary Figure 16D*).

In the testis, our gene expression profiling of BM and NBMs revealed that there are no obvious differences, unlike what was observed between Q vs NBF ovaries. This could be partly explained by the fact that NBFs do not produce mature gametes but Q’s do, whereas both BMs and NBMs NMRs undergo spermatogenesis to produce gametes, and this is reflected in spermatogenesis gene expression profiles seen in both. Nevertheless, some DEGs were identified that are known to play an important role in male sexual differentiation, reproduction, and responses to steroids. In particular, the enrichment of GO term involved in male reproduction (meiotic cell cycle, male genitalia development, gamete generation, spermatogenesis, and sperm motility) for upregulated DEGs in BMs includes genes such as *Prm1, Prm2, Odf3 and Akap4*, which have been shown to have a important role in mouse spermatogenesis. A decrease in expression of genes such as *Prm1, Prm2, Odf3* and *Akap4* and related genes in NBMs, might contribute for the observed reduction in sperm number and motility [28]. This is further complemented by the low expression of genes that are involved in meiotic and post-meiotic stages of spermatogenesis in non-breeding animals. Based on these observations, we suggest that the main difference between BMs and NBMs spermatogenesis is related to post-meiotic and sperm maturation stages (*Supplementary Figure 3E*), where NBMs fail to express critical genes at appropriate level at these stages (post-meiotic and sperm maturation stages), resulting in low sperm count and impaired motility. Cytological examination of testicular sections of breeding and non-breeding animals (*Supplementary Figure 15*), confirms previous observations of a higher interstitial cell content in breeding animals compared to non-breeding animals. At the gene expression level, we also observed a significant decrease (2 fold, *Additional file 29*) in the expression of the LH receptor gene (*Lhcgr*) in NBMs compared to BMs. This fits with predictions based on the reduction in interstitial (Leydig) cells numbers, together with previously reported observations of lower concentrations of urinary testosterone and plasma LH in NBMs [20].

## Conclusion

Our study reveals gene expression differences among male and female NMR reproductive castes. Our findings in brain, ovaries and testes provide some of the first insights into the potential molecular mechanisms that are important in reproduction suppression of NMR. The gene expression differences in the NMR brains follow social status rather than sex, with the Q having a unique gene expression pattern compared to the rest of the colony. In highlighting the potential importance of dopamine pathways in the brain, and a possible role for hyperprolactinemia in mediating suppression, our findings significantly advance the understanding of the basis of mammalian eusociality and cooperative breeding systems. In the ovaries, Qs express genes that are required for estrogen production, and thereby undergoes oogenesis/folliculogenesis, ovulation and other sexual differentiation processes. In contrast, NBFs are arrested at a pre-pubertal stage, and do not have the ability to produce ovarian estrogen, explaining their failure to ovulate and subsequent reproductive incompetency. In the testis, both BMs and NBMs undergo spermatogenesis, however NBMs fail to express genes required for post-meiotic sperm maturation, giving explanation to lower sperm numbers, impaired motility, and breeding incapacity of NBMs. While many questions remain, the phenotypic plasticity exhibited by the NMR also offers scope for understanding the dynamics of reproductive activation when non-breeders transition into breeding state (NBF to Qs or NBM to BM), in particular the role of epigenetics and other regulatory factors affecting the genes associated in this transformation.

## Material and methods

### Animals

NMRs were maintained at Queen Mary University of London in compliance with institutional guidelines. All animals were born in captivity, kept under constant ambient temperature of 28-30°C, and housed in artificial burrow systems composed of interconnected perspex tubing with separate chambers for nesting, food and toilet, simulating natural burrow conditions. They were fed an ad libitum diet of a variety of chopped root vegetables such as sweet potato and turnip. Animal tissues were collected post mortem (*Additional file 1A*) following euthanasia in full accordance with Institutional and National animal care and use guidelines.

### Blood sampling for hormone assay

Blood samples were obtained from *X NBF, Queen BM, NBM naked mole-rats* from among 13 captive colonies from QMUL and UP. All blood samples collected at the University of Pretoria were with local ethics committee clearance. Blood samples were collected between 11h00 and 15h00 as follows: The animals were hand held and venous blood samples collected from the hind foot. Approximately 300-500ul of blood was collected into heparinised micro-haematocrit tubes (University of Pretoria samples) or into a heparinised 1 ml syringe from trunk blood following euthanasia (QMUL) prior to tissue collection. The blood was centrifuged at 500g for 15 minutes and the plasma separated from the red cells and stored at −80ºC until hormone assay.

### RNA extraction and sequencing

NMR brain (excluding the olfactory bulb, the cerebellum, the medulla and the pons), ovary, and testis were snap frozen by immersion into liquid nitrogen and subsequently stored at −80ºC prior to RNA extraction. Snap frozen tissues were mechanically powdered and mixed to maintain the heterogeneity of the sample. Total RNA was extracted from tissues using Qiagen miRNeasy kit (Qiagen, USA) following the manufacturer’s recommendations. The quality of the extracted RNA was controlled using the Agilent Bioanalyzer (Agilent) and Qubit Fluorometric Quantitation (Thermo Fisher Scientific Inc.). 1µg of high quality RNA (with RNA Integrity Number (RIN) >7) were used for RNA sequencing. Total RNA, after polyadenylated RNA purification, was prepared for sequencing using Illumina Truseq library preparation protocol. For each sample, around 100 (*Additional file 1B*) million raw paired-end sequence reads (101 base pair long) were generated using Illumina HiSeq 2000 sequencing instrument. Data sets are available from NCBI GEO under accession number (####).

### RNA sequence (RNA-seq) analysis

The quality of the generated RNA sequence was evaluated using FastQC (version 0.11.2) [119]. Sequence adapters, low quality reads, and overrepresented sequences (e.g. mitochondrial sequences) in the reads were removed using Trimmomatic (version 0.32) [120], and the quality of the reads was checked again using FastQC. Sequence that passed the quality assessment were aligned to the NMR genome (hetGla2, Broad institute, 2012) using tophat2 (version 2.0.6, default parameters) [121], with bowtie2 (version 2.1.0) [122]. For each sample genome guided de-novo transcriptome assembly was performed using Cufflinks (version 2.2.1, default parameters) [123] and assembled transcript from all samples were merged using cuffmerge (cufflinks) to generate a master reference transcripts. Merged transcripts (master transcripts) were annotated to gene name using mouse transcripts. Mouse cDNA were obtained from Ensembl [124], and a mouse cDNA blast database was generated using blast (version 2.2.25) [125, 126]. Assembled and merged NMR transcripts (master transcript) were searched against mouse cDNA blast database, and transcripts with e val <= 10∧-5, with a length of >200bp were retained. Transcript abundance level was generated using master reference transcripts generated by cufflinks and HTSeq (version 0.5.3p9) [127]. The transcript level quantified using HTSeq was used as an input for further processing using R software environment for statistical computing and graphics (version 3.2.2). Data normalization, removal of batch effect and other variant was performed using EDASeq R package (version 2.2.0) [128] and RUVseq package (Remove Unwanted Variation from RNA-Seq package) [129]. In short, read counts were normalized using EDASeq R package, and “in-silico empirical” negative controls genes, for RUVseq package (RUVg normalization) were obtained by taking least significantly DEGs based on a first-pass differential expression analysis performed prior to RUVg normalization. RUVseq package (RUVg normalization) was then performed using the “in-silico empirical” negative controls genes. Differential expression was performed using edgeR R package (version 3.10.5) [130], using the negative binomial GLM approach, edgeR normalization on raw counts, and by considering a design matrix that includes both the covariates of interest and the factors of unwanted variation. DEGs with false discovery rate (FDR <=0.05, Benjamini-Hochberg multiple testing correction), expression level > 1 CPM (counts per million), and log fold change >1 were retained and used for further processing, gene ontology and pathway analysis.

### Gene ontology and pathway analysis

Gene ontology and pathway analysis were performed using Metascape (metascape.org) [131], Enrichr [132], and Ingenuity Pathway Analysis (IPA^®^, QIAGEN Redwood City, www.qiagen.com/ingenuity). In Metascape, for each given gene list of DEGs, pathway and process enrichment analysis was carried out with the following ontology sources: GO Biological Processes, GO Molecular Functions and KEGG Pathway. All genes in the genome were used as the enrichment background. Terms with p-value < 0.01, minimum count 3, and enrichment factor > 1.5 (enrichment factor is the ratio between observed count and the count expected by chance) are collected and grouped into clusters based on their membership similarities [131]. P-values are calculated based on accumulative hypergeometric distribution, q-values are calculated using the Benjamini-Hochberg procedure to account for multiple testing [131]. Kappa scores were used as the similarity metric when performing hierarchical clustering on the enriched terms and then sub-trees with similarity > 0.3 are considered a cluster. The most statistically significant term within a cluster is chosen as the one representing the cluster [131]. To further capture the relationship among terms, a subset of enriched terms were selected and rendered as a network plot, where terms with similarity > 0.3 are connected by edges, with the best p-values from each of the 20 clusters [131]. Similar independent enrichment analysis was performed using Enrichr [132], to validate the outcome of Metascape. The list of DEGs were used as input to Enrichr and the enrichment of GO Biological Processes, GO Molecular Functions was investigated. The final result was sorted and plotted by using a combined score. The combined score is a combination of the p-value and z-score calculated by multiplying the two scores as follows: c = log(p) * z, where c is the combined score, p is the p-value computed using Fisher’s exact test, and z is the z-score computed to assess the deviation from the expected rank [132].

### Tissue preparation and immunofluorescence

NMRs were deeply anesthetized with 80 mg pentobarbital (400 µL Euthatal, IP) and transcardially perfused with 40 g/L formaldehyde in PBS for 10 min (10 mL/min). Brains were dissected and kept at 4°C for 12h in a PBS solution containing 15% sucrose before being flash frozen in isopentane (1 min at − 30°C) and stored at −80°C. Serial coronal sections (30 µm thick) were made with a cryostat (Leica, France) and mounted onto Superfrost Plus slides. For each animal, the first section containing some frontal cortex was taken as the origin of the numbering. Sections were stored dry at −80°C until being processed for immunolabelling.

Sections were equilibrated to −20°C before a short additional fixation with 30 g/L formaldehyde in PBS for 5 min at RT. Formaldehyde was then neutralized with TBS (50 mM Tris, 150 mM NaCl, pH 7.4) for 5 min at 4°C, before two consecutive steps of permeabilization of 5 min each at 4°C, first in PBS containing 0.5% (vol/vol) Triton X-100, then in PBS with 10% (vol/vol) methanol. The sections were rinsed with 70% (vol/vol) ethanol (EtOH70) and kept in the same solution for 10 min at RT before being treated with 0.1% (w/vol) Sudan Black B in EtOH70, for autofluorescence removal. After two quick rinses with EtOH70 and one PBS wash, 30 min preblocking at 4°C was achieved with the IF buffer ie PBS, 2% (w/vol) BSA, 0.2% (vol/vol) Tween 20, 50 mM glycine. The anti-TH rabbit polyclonal antibody (Abcam, ab112) diluted 1/800 in IF buffer was incubated overnight at 4°C. Then the sections were rinsed 3 times 10 minutes with PBS before being incubated with an Alexa488-coupled goat anti-rabbit secondary antibody (Invitrogen A-11034) diluted 1/500 in PBS for 1h at RT. After 3 rinses in PBS and a 30 min DAPI staining step (100 nM in PBS), sections were finally rinsed and mounted in Vectashield (Vector Laboratories, USA). Image acquisition was carried out at the Cell and Tissue Imaging Platform of the Genetics and Developmental Biology Department (UMR3215/U934) of Institut Curie. The sections of the different animals were processed at the same time with the same parameters. Full views of the sections consisted of mosaics of pictures made with the 5X objective of an upright epifluorescence microscope (Zeiss). Views of the hippocampus were made with the 10X objective. Additional z-stack pictures (1 µm steps) of the regions of interest were taken for quantification, with the 10X objective of an upright spinning disk confocal microscope (Roper/Zeiss). For each picture, 5 consecutive confocal optical sections were summed, then the average intensity of the region of interest was measured, and finally the measures for both hemispheres were averaged (when applicable).

### Histology of testes and ovaries

Gonadal tissue samples were fixed by immersion in 4% paraformaldehyde in saline within 15 minutes of collection for a minimum of seven days. After fixation tissue samples were dehydrated and cleared using standard histological methodology, before embedding in paraffin wax. Sections of 5-8 µm were cut and stained for light microscopy with haematoxylin-eosin. Photomicrographs of sections were captured with a QIClick™ CCD Camera (01-QICLICK-R-F-CLR-12; QImaging) linked to a DMRA2 light microscope (Leica), using Volocity^®^ v.6.3.1 image analysis software (Perkin-Elmer) running on an iMac computer (27-inch with OS X Yosemite, version 10.10).

### Prolactin assay and validation

Plasma prolactin concentrations were determined using a commercial enzyme-linked immunosorbent assay (Elabscience© Guinea pig prolactin ELISA kit, Catalogue No: E-EL-GP0358) according to the instructions in the manufacturer’s user manual. In brief, 100 ml of reference standard and diluted samples (1/2 to 1/50 in sample diluent) were transferred into coated wells of a 96-well micro-ELISA plate, respectively, and incubated for 90 min at 37°C. Subsequently, all supernatant was removed, and the plate patted dry. Immediately, 100 ml of biotinylated detection antibody was added, and incubated for 60 min at 37°C. The plate was washed 3 times, patted dry, and 100 ml of horse radish peroxide (HRP) conjugate were added and incubated for 30 min at 37°C. Subsequently, the plate was washed 5 times with washbuffer, and patted dry. 90 ml of substrate reagent were added, and incubated for 15 min at 37°C. To terminate the enzymatic reaction, 50 ml of stop solution were added. Optical density was determined at 450 nm and results calculated using a best-fit curve. The sensitivity of the assay was 0.1 ng/ml, the detection range 0.16-20 ng/ml, and coefficient of variation for repeatability was < 10%.

## Acknowledgments

The authors would like to acknowledge the Cell and Tissue Imaging Platform of the Genetics and Developmental Biology Department (UMR3215/U934) of *Institut Curie*, member of France-Bioimaging (supported by ANR-10-INSB-04), particularly Olivier Renaud for help with light microscopy. We thank all members of the Edith Heard team for stimulating discussions, technical and conceptual advice, notably Mikael Attia and Ronan Chaligné. We thank Deborah Bourc’his, Nicolas Servant, and Emmanuel Barillot from Institut Curie for their insightful comments on the manuscript. We would like to acknowledge the Biological Services Unit staff at QMUL (Richard Rountree, Arthur Gatward, Paul Sroka) for care and maintenance of animals and Cécile Reyes (Institut Curie, Genomics facility) for tissue fragmentation and RNA extraction, and Professor Jenny Jarvis (University of Cape Town). We thank the Institut Curie High-throughput sequencing platform (NGS platform) for generating all the sequencing data.

## Availability of data and materials

The datasets generated and/or analysed during the current study are available in the NCBI GEO repository (https://www.ncbi.nlm.nih.gov/geo/), under series number #####

## Author contributions

EM conceived the study, designed experimental strategies, performed all bioinformatic analyses, interpreted results, performed experiments and wrote the manuscript. LMP performed and interpreted brain validation experiments, and participated in writing the manuscript. DG performed RNA extraction. NCB collected plasma samples and funded hormonal work, while SBG and AG carried out hormone assays. EHB designed experimental strategies, interpreted results, funded the project. CGF conceived the study, designed experimental strategies, interpreted results, performed experiments and wrote the manuscript. EH conceived the study, designed experimental strategies, interpreted results, and wrote the manuscript.

## Ethical approval

Not applicable

## Consent for publication

Not applicable

## List of abbreviations

NMR: Naked mole-rats
LH: Luteinizing hormone
GnRH: Gonadotrophin releasing hormone
RNA-seq: RNA-sequencing
PCA: Principal component analysis
Q: Queen
Qs_Tr: Queen technical replicate
NBF: Non-breeding females
NBF_Tr: Non breeding female technical replicate
NBM: Non-breeding males
BM: Breeding males
CPM: Counts per million
DEG: Differentially expressed genes
GO: Gene ontology
Q: genes Queen genes
TH/Th: Tyrosine hydroxylase
FSH: Follicle-stimulating hormone
PGCs: Primordial germ cells
PRL: Prolactin
bp: Base pair
IF: Immunofluorescene
PValue: p-value
logCPM: Counts per million in log scale
logFC: Log fold change
FDR: False discovery rate
BH: Benjamini-Hochberg

## Supplementary Figure legends

**Supplementary Figure 1: A)** Principal component analysis (PCA) plots showing the clustering of different NMR colony members based on global brain gene expression profile. **B)** Cluster dendrogram showing hierarchical clustering of the NMR brain gene expression. Hierarchical clustering was generated using euclidean distance matrix computation and ward.D2 agglomeration method. **C)** Enriched biological process terms for DEGs that are common in the comparison between Qs vs NBMs and Qs vs NBMs (Q genes) generated using Enrichr. The length of the bar represents the significance of that specific gene-set or term, and the color intensity provides additional information about the significance (the brighter the color, the more significant that term is). Statistical information used to generate the graph including the p-value and other enriched terms are provided in *Additional file 18*. **D)** Enriched molecular functions terms for DEGs that are common in the comparison between Qs vs NBMs and Qs vs NBMs (Q genes) generated using Enrichr. The length of the bar represents the significance of that specific gene-set or term, and the color intensity provides additional information about the significance (the brighter the color, the more significant that term is). Statistical information used to generate the graph including the p-value and other enriched terms are provided in *Additional file 19*.

**Supplementary Figure 2: Gene expression differences between the brains of Qs and NBFs. A)** Principal component analysis (PCA) plot showing the clustering of Qs and NBFs brains based global gene expression profile. **B)** Cluster heatmap of Qs and NBFs brain gene expression. Sample distance was calculated euclidean distance matrix computation and cluster agglomeration was done using ward.D2 method. Heatmap color indicate the euclidean distance between samples indicated in the heatmap color key. **C)** Volcano plot showing significance versus fold-change. The log fold change in expression is indicated on the x-axis and significance -log_10_(p-value) is indicated on the y-axis. DEGs are indicated by red color. **D)** Heatmap of enriched terms (Canonical Pathways, GO Biological Processes, Hallmark Gene Sets, KEGG Pathway) for DEGs between Qs and NBFs (generated using Metascape). Significance of enrichment is indicated on the x-axis in –log_10_(p-value). The color code on the y-axis is used to show the clustering and relation of these networks (shown in *Supplementary Figure 2E*). Detailed list of enriched terms can be found at *Additional file 3*. **E)** Network of enriched terms (*Supplementary Figure 2D*) colored by cluster ID, indicating the relationship between the different enriched terms (cluster ID color correspond to the color code shown in *Supplementary Figure 2D* y-axis). Nodes that share the same cluster are close to each other. Each circle node represents a term and the size of the circle is proportional to the number of genes that fall into that term, and the identity of the cluster is indicated by its color (nodes of the same color belong to the same cluster). Similar terms are linked by an edge (the thickness of the edge represents the similarity score). One term from each cluster is selected to have its term description shown as label. **E)** Enriched biological process terms for DEGs that are common in the comparison between Qs vs NBFs generated using Enrichr. The length of the bar represents the significance of that specific gene-set or term, and the color intensity provides additional information about the significance (the brighter the color, the more significant that term is). Statistical information used to generate the graph including the p-value and other enriched terms are provided in *Additional file 4*. **G)** Enriched molecular functions terms in for DEGs that are common in the comparison between Qs vs NBFs generated using Enrichr. The length of the bar represents the significance of that specific gene-set or term, and the color intensity provides additional information about the significance (the brighter the color, the more significant that term is). Statistical information used to generate the graph including the p-value and other enriched terms are provided in *Additional file 5*.

**Supplementary Figure 3: Gene expression differences between the brains of BMs NBMs. A)** Scatter plot of testis size to body weight. The size of the circles in the scatter plot are proportional to testis to body weight ratio. BMs are depicted in blue and NBMs in black. **B)** Principal component analysis (PCA) plot showing the clustering of BMs and NBMs based global brain gene expression profile. **C)** Cluster heatmap of BMs and NBMs brain gene expression. Sample distance was calculated euclidean distance matrix computation and cluster agglomeration was done using ward.D2 method. Heatmap colors indicate the euclidean distance between samples indicated in the heatmap color key. **D)** Volcano plot showing the significance versus fold-change. The log fold change in expression in indicated on the x-axis and significance in -log_10_(p-value) is indicated on the y-axis. DEGs are indicated by the red color. **E)** Heatmap of enriched terms (Canonical Pathways, GO Biological Processes, Hallmark Gene Sets, KEGG Pathway) for DEGs between BMs and NBMs (generated using Metascape). Significance of enrichment is indicated on the x-axis in –log_10_(p-value). The color code on the y-axis is used to show the clustering and relation of this networks (shown in *Supplementary Figure 3F*). Detailed list of enriched terms can be found at *Additional file 7*. **F)** Network of enriched terms (*Supplementary Figure 3E*) colored by cluster ID, indicating the relationship between the different enriched terms (cluster ID color correspond to the color code shown in *Supplementary Figure 3E y-axis*). Nodes that share the same cluster are close to each other. Each circle node represents a term and the size of the circle is proportional to the number of genes that fall into that term, and the identity of the cluster is indicated by its color (nodes of the same color belong to the same cluster). Similar terms are linked by an edge (the thickness of the edge represents the similarity score). One term from each cluster is selected to have its term description shown as label. **G)** Enriched biological process terms for DEGs that are common in the comparison between BMs vs NBMs generated using Enrichr. The length of the bar represents the significance of that specific gene-set or term, and the color intensity provides additional information about the significance (the brighter the color, the more significant that term is). Statistical information used to generate the graph including the p-value and other enriched terms are provided in *Additional file 8*. **H)** Enriched molecular functions terms in for DEGs that are common in the comparison between BMs vs NBMs generated using Enrichr. The length of the bar represents the significance of that specific gene-set or term, and the color intensity provides additional information about the significance (the brighter the color, the more significant that term is). Statistical information used to generate the graph including the p-value and other enriched terms are provided in *Additional file 9*.

**Supplementary Figure 4: Gene expression differences between the brains of Qs and BMs. A)** Principal component analysis (PCA) plot showing the clustering of Qs and BMs based global brain gene expression profile. **B)** Cluster heatmap of Qs and BMs brain gene expression. Sample distance was calculated euclidean distance matrix computation and cluster agglomeration was done using ward.D2 method. Heatmap color indicates the euclidean distance between samples indicated in the heatmap color key. **C)** Volcano plot showing significance versus fold-change. The log fold change in expression is indicated on the x-axis and significance in -log_10_(p-value) is indicated on the y-axis. DEGs are indicted by the red color. **D)** Heatmap of enriched terms (Canonical Pathways, GO Biological Processes, Hallmark Gene Sets, KEGG Pathway) for DEGs between Qs and BMs (generated using Metascape). Significance of enrichment is indicated on the x-in –log_10_(p-value). The color code on the y-axis is used to show the clustering and relation of this networks (shown in *Supplementary Figure 4E*). Detailed list of enriched terms can be found at *Additional file 11*. **E)** Network of enriched terms (*Supplementary Figure 4D*) colored by cluster ID, indicating the relationship between the different enriched terms (cluster ID color correspond to the color code shown in *Supplementary Figure 4D y-axis*). Nodes that share the same cluster are close to each other. Each circle node represents a term and the size of the circle is proportional to the number of genes that fall into that term, and the identity of the cluster is indicated by its color (nodes of the same color belong to the same cluster). Similar terms are linked by an edge (the thickness of the edge represents the similarity score). One term from each cluster is selected to have its term description shown as label. **F)** Enriched biological process terms for DEGs that are common in the comparison between Qs vs BMs generated using Enrichr. The length of the bar represents the significance of that specific gene-set or term, and the color intensity provides additional information about the significance (the brighter the color, the more significant that term is). Statistical information used to generate the graph including the p-value and other enriched terms are provided in *Additional file 12*. **G)** Enriched molecular functions terms for DEGs that are common in the comparison between Qs vs BMs generated using Enrichr. The length of the bar represents the significance of that specific gene-set or term, and the color intensity provides additional information about the significance (the brighter the color, the more significant that term is). Statistical information used to generate the graph including the p-value and other enriched terms are provided in *Additional file 13*.

**Supplementary Figure 5: Gene expression differences between the brains of NBFs and NBMs. A)** Cluster heatmap of NBFs and NBMs gene brain gene expression. Sample distance was calculated euclidean distance matrix computation and cluster agglomeration was done using ward.D2 method. Heatmap color indicate the euclidean distance between samples indicated in the heatmap color key. **B)** Volcano plot showing the significance versus fold-change. The log fold change in expression in indicated on the x-axis and significance in -log_10_(p-value) is indicated on the y-axis. DEGs are indicated by the red color.

**Supplementary Figure 6: Venn diagrams showing the number of DEGs in the brain that were identified in comparisons among different sex/status groups, and the relationship between them. A)** Venn diagram showing all DEGs that are common in the comparison between Qs vs NBFs, Qs vs NBMs, and Qs vs BMs. **B)** Venn diagram showing DEGs that also shows higher expression in the comparison between Qs vs NBFs, Qs vs NBMs, and Qs vs BMs. **C)** Venn diagram showing DEGs that also show lower expression in the comparison between Qs vs NBFs, Qs vs NBMs, and Qs vs BMs. **D)** Venn diagram showing DEGs that also show higher expression in the comparison between Qs vs NBFs and Qs vs NBMs. **E)** Venn diagram showing DEGs that also show lower expression in the comparison between Qs vs NBFs and Qs vs NBMs. **F)** Venn diagram showing all DEGs that are common in the comparison between BMs vs NBFs and BMs vs NBMs. **G)** Venn diagram showing DEGs that also show higher expression in the comparison between BMs vs NBFs and BMs vs NBMs. **H)** Venn diagram showing DEGs that also show lower expression in the comparison between BMs vs NBFs and BMs vs NBMs.

**Supplementary Figures 7: Gene expression profile of Q and NBF ovaries. A)** Cluster heatmap of ovary samples. Sample distance was calculated euclidean distance matrix computation and cluster agglomeration was done using ward.D2 method. Heatmap color indicates the euclidean distance between samples indicated in the heatmap color key. **B)** Volcano plot showing the significance versus fold-change. The log fold change in expression in indicated on the x-axis and significance in -log_10_(p-value) is indicated on the y-axis. DEGs are indicated by the red color. **C)** Network of enriched terms colored by cluster ID, indicating the relationship between the different enriched terms (*Figure 3D*). Nodes that share the same cluster are close to each other. Each circle node represents a term and the size of the circle is proportional to the number of genes that fall into that term, and the identity of the cluster is indicated by its color (nodes of the same color belong to the same cluster). Similar terms are linked by an edge (the thickness of the edge represents the similarity score). One term from each cluster is selected to have its term description shown as label. **D)** Enriched biological processes for DEGs between Qs vs NBFs ovary generated using Enrichr. The length of the bar represents the significance of that specific gene-set or term, and the color intensity provides additional information about the significance (the brighter the color, the more significant that term is). Statistical information used to generate the graph including the p-value and other enriched terms are provided in *Additional file 27a*. **E)** Enriched molecular functions for DEGs between Qs vs NBFs ovary generated using Enrichr. The length of the bar represents the significance of that specific gene-set or term, and the color intensity provides additional information about the significance (the brighter the color, the more significant that term is). Statistical information used to generate the graph including the p-value and other enriched terms are provided in *Additional file 28A*.

**Supplementary Figure 8: Gene enrichment for DEGs that also show lower expression in Q ovaries. A)** Heatmap of enriched terms (Canonical Pathways, GO Biological Processes, Hallmark Gene Sets, KEGG Pathway) for DEGs that show lower expression in Q ovary (generated using Metascape). The color of the bar graph is proportion to the p-values. Significance of enrichment is indicated on the x-axis in –log_10_(p-value). The color code on the y-axis is used to show the clustering and relation of these networks (shown in *Supplementary Figure 8B*). More information about the enriched terms can be found in *Additional file 26B*. **B)** Network of enriched terms colored by cluster ID, indicating the relationship between the different enriched terms (*Supplementary Figure 8A*). Nodes that share the same cluster are close to each other. Each circle node represents a term and the size of the circle is proportional to the number of genes that fall into that term, and the identity of the cluster is indicated by its color (nodes of the same color belong to the same cluster). Similar terms are linked by an edge (the thickness of the edge represents the similarity score). One term from each cluster is selected to have its term description shown as label. **C)** Enriched biological processes for DEGs between Qs and NBFs ovary and also show lower expression in the Qs generated using Enrichr. The length of the bar represents the significance of that specific gene-set or term, and the color intensity provides additional information about the significance (the brighter the color, the more significant that term is). Statistical information used to generate the graph including the p-value and other enriched terms are provided in *Additional file 27B*. **D)** Enriched molecular functions for DEGs between Qs and NBFs ovary and also show lower expression in the Qs generated using Enrichr. The length of the bar represents the significance of that specific gene-set or term, and the color intensity provides additional information about the significance (the brighter the color, the more significant that term is). Statistical information used to generate the graph including the p-value and other enriched terms are provided in *Additional file 28B*.

**Supplementary Figure 9: Gene enrichment for DEGs that also show higher expression in Q ovaries. A)** Heatmap of enriched terms (Canonical Pathways, GO Biological Processes, Hallmark Gene Sets, KEGG Pathway) for DEGs that show higher expression in Qs ovary (generated using Metascape). The color of the bar graph is proportion to the p-values. Significance of enrichment is indicated on the x axis in –log_10_(p-value). The color code on the y-axis is used to show the clustering and relation of this networks (shown in *Supplementary Figure 9B*). More information about the enriched terms can be found in *Additional file 26C*. **B)** Network of enriched terms colored by cluster ID, indicating the relationship between the different enriched terms (*Supplementary Figure 9A*). Nodes that share the same cluster are close to each other. Each circle node represents a term and the size of the circle is proportional to the number of genes that fall into that term, and the identity of the cluster is indicated by its color (nodes of the same color belong to the same cluster). Similar terms are linked by an edge (the thickness of the edge represents the similarity score). One term from each cluster is selected to have its term description shown as label. **C)** Enriched biological processes for DEGs between Qs and NBFs ovary and also show higher expression in the Qs generated using Enrichr. The length of the bar represents the significance of that specific gene-set or term, and the color intensity provides additional information about the significance (the brighter the color, the more significant that term is). Statistical information used to generate the graph including the p-value and other enriched terms are provided in *Additional file 27C*. **D)** Enriched molecular functions for DEGs between Qs and NBFs ovary and also show higher expression in the Q generated using Enrichr. The length of the bar represents the significance of that specific gene-set or term, and the color intensity provides additional information about the significance (the brighter the color, the more significant that term is). Statistical information used to generate the graph including the p-value and other enriched terms are provided in *Additional file 28C*.

**Supplementary Figure 10: NMR Q and NBF ovary histology.** Representative sections (at the same magnification) through the ovary of non-breeding (**A**) and Queen (**B**) NMRs: S stroma, O oocyte, P primordial follicle, 1º primary follicle, 2º secondary follicle, 2º* secondary follicle with two oocytes, 3º tertiary follicle.

**Supplementary Figure 11: Gene expression levels of NMR genes at different stages of mouse oogenesis/folliculogenesis. A**) Violin plot combined with boxplot showing the expression level of NMR Qs and NBFs (ovary RNA-seq data) for clusters that show stage specific expression during mouse oogenesis/folliculogenesis. Mouse gene cluster that show a decrease in expression from the primary to small antral stage (cluster 1) and another gene cluster (cluster 5: where gene expression increased at small antral and large antral follicle follicle stages) were obtained from (50) (753 genes in total in the two clusters). NMR Qs and NBFs ovary expression level was mapped to these mouse folliculogenesis genes names (cluster 1 and cluster 5), and the expression level of these genes in the Q and NBF NMR ovaries was plotted (x-axis: clusters, y-axis: expression level in log_2_). **B)** Violin plot in combination with boxplot showing the expression level of Qs and NBFs genes (NMR ovary) for DEGs that show up and down regulation in mouse infant vs adult whole ovary comparison (51). From 7021 DEGs that show up or down regulation in mouse infant vs adult whole ovary gene expression comparison (51) 4494 genes that show >2 fold change (up or down) were extracted, and their expression in Qs and NBFs (NMR ovary) was plotted. x-axis: differential expression group (down, DEGs that show lower expression in adult mouse ovary compared to infant; and up, for genes that show higher expression in adult mouse ovary compared to infant); y-axis expression level in log_2_ scale. **C)** Bar plot showing the expression level of mouse oocyte specific genes in NMR Qs and NBFs ovary RNA-seq data. Mouse oocyte specific genes were obtained from [ref], and their average expression level in Q and NBF NMR ovaries (RNA-seq data) was plotted (x-axis: gene name, y-axis: expression level in log_2_ scale). **D)** Expression of *Cyp19a1* (*Aromatase*) in Q and NBF NMR ovaries (x-axis, sample name; y-axis, expression level in counts per million (CPM)).

**Supplementary Figure 12**: **Gene expression profile of BMs and NBMs testis, and enriched terms for DEGs. A)** Cluster heatmap of testis samples. Sample distance was calculated euclidean distance matrix computation and cluster agglomeration was done using ward.D2 method. Heatmap colors indicate the euclidean distance between samples shown in the heatmap color key. **B)** Volcano plot showing the significance versus fold-change. The log fold change in expression in indicated on the x-axis and significance in -log_10_(p-value) is indicated on the y-axis. DEGs are indicated by the red color. **C)** Network of enriched terms colored by cluster ID, indicating the relationship between the different enriched terms (*Figure 4F*). Nodes that share the same cluster are close to each other. Each circle node represents a term and the size of the circle is proportional to the number of genes that fall into that term, and the identity of the cluster is indicated by its color (nodes of the same color belong to the same cluster). Similar terms are linked by an edge (the thickness of the edge represents the similarity score). One term from each cluster is selected to have its term description shown as label. **D)** Enriched biological processes (generated using Enrichr). The length of the bar represents the significance of that specific gene-set or term, and the color intensity provides additional information about the significance (the brighter the color, the more significant that term is). Statistical information used to generate the graph including the p-value and other enriched terms are provided in *Additional file 31A*. **E)** Enriched molecular functions (generated using Enrichr). The length of the bar represents the significance of that specific gene-set or term, and the color intensity provides additional information about the significance (the brighter the color, the more significant that term is). Statistical information used to generate the graph including the p-value and other enriched terms are provided in *Additional file 32A*.

**Supplementary Figure 13**: **Gene enrichment for DEGs that also show lower expression in BM testes. A)** Heatmap of enriched terms (Canonical Pathways, GO Biological Processes, Hallmark Gene Sets, KEGG Pathway) for DEGs that show lower expression in BMs (generated using Metascape). The color code on the y-axis is used to show the clustering and relation of enriched networks (*Supplementary Figure 13B*). Detailed list of enriched terms and genes that belong to these terms is provided in *Additional file 30B*. **B)** Network of enriched terms found in above colored by cluster ID, indicating the relationship between the different enriched terms (*Supplementary Figure 13A*). Gene enrichment was generated using DEGs that show higher expression in BMs compared to NBMs. Nodes that share the same cluster are close to each other. Nodes that share the same cluster are close to each other. Each term is represented by a circle node, where its size is proportional to the number of input genes fall into that term, and its color represent its cluster identity (i.e., nodes of the same color belong to the same cluster). Similar terms are linked by an edge (the thickness of the edge represents the similarity score). One term from each cluster is selected to have its term description shown as label. **C)** Enriched biological processes (generated using Enrichr). The length of the bar represents the significance of that specific gene-set or term, and the color intensity provides additional information about the significance (the brighter the color, the more significant that term is). Statistical information used to generate the graph including the p-value and other enriched terms are provided in *Additional file 31B*. **D)** Enriched molecular functions (generated using Enrichr). The length of the bar represents the significance of that specific gene-set or term, and the color intensity provides additional information about the significance (the brighter the color, the more significant that term is). Statistical information used to generate the graph including the p-value and other enriched terms are provided in *Additional file 32B*.

**Supplementary Figure 14**: **Gene enrichment for DEGs that also show higher expression in breeding male testes. A)** Heatmap of enriched terms (Canonical Pathways, GO Biological Processes, Hallmark Gene Sets, KEGG Pathway) for DEGs that show higher expression in BMs (generated using Metascape). Enriched terms significance is colored by p-values. The color code on the y-axis is used to show the clustering and relation of enriched networks (*Supplementary Figure 14B*). Detailed list of enriched terms and genes that belong to these terms is provided in *Additional file 30C*. **B)** Network of enriched terms found in above colored by cluster ID, indicating the relationship between the different enriched terms (*Supplementary Figure 14A*). Gene enrichment was generated using genes that show higher expression in breeding males compared to NBMs. Nodes that share the same cluster are close to each other. Nodes that share the same cluster are close to each other. Each term is represented by a circle node, where its size is proportional to the number of input genes fall into that term, and its color represent its cluster identity (i.e., nodes of the same color belong to the same cluster). Similar terms are linked by an edge (the thickness of the edge represents the similarity score). One term from each cluster is selected to have its term description shown as label. **C)** Enriched biological processes. The length of the bar represents the significance of that specific gene-set or term, and the color intensity provides additional information about the significance (the brighter the color, the more significant that term is). Statistical information used to generate the graph including the p-value and other enriched terms are provided in *Additional file 31C*. **D)** Enriched molecular functions. The length of the bar represents the significance of that specific gene-set or term, and the color intensity provides additional information about the significance (the brighter the color, the more significant that term is). Statistical information used to generate the graph including the p-value and other enriched terms are provided in *Additional file 32C*.

**Supplementary Figure 15: NMR BM and NBM testis histology.** Representative sections through the testes of adult (**A,C**) breeding and (**B,D**) non-breeding male NMRs showing spermatogenesis; (**A**) and (**B**) are higher magnification showing examples of the cell types at different stages: S_A_ spermatogonia Type A, S_B_ spermatogonia Type B, S_1_ primary spermatocyte, S_3_ spermatid, S_4_ spermatocyte, S_t_ Sertoli cell (S_2_ secondary spermatocytes not identified). Sections (**C**) and (**D**) are lower magnification (from the respective individuals) and show the marked differences in the relative amount of interstitial tissue (mainly Leydig cells) to seminiferous tubules (bounded by the dotted lines).

**Supplementary Figure 16: A)** Potentially dopaminergic neuron cell groups (A8-A16), expressing the enzyme tyrosine hydroxylase but not the dopamine beta-hydroxylase, with their principal projections in the adult rodent brain (adapted from (45)). **B)** and **C)** A model describing the possible pathways though which a dopamine could regulate reproductive division of labor; **B)** in breeding animals (Qs and BMs), increased dopamine production in the hypothalamus suppresses the production of prolactin in from the pituitary. In the absence (or low amounts) of prolactin, GnRH is released from the hypothalamus, which acts on anterior pituitary to facilitate the release of FSH and LH. FSH and LH act on the gonads of breeding animals (Qs and BMs), to bring about normal gonadal development, gametogenesis and the release of sex hormones (estrogen and testosterone). **C)** in non-breeding naked mole-rats animals, lower levels of dopamine release result in increased production of prolactin by the anterior pituitary, which in turn suppresses the release of GnRH from the hypothalamus. In the absence of normal GnRH secretion, the anterior pituitary produces low levels of FSH and LH, resulting in suppression of normal gonadal development, gametogenesis and the release of sex hormones. **D)** A model showing the possible arrest point for NBF NMRs (indicated by the color graduation in the horizontal bar). The Q ovary can complete all the stages of oogesisis, however NBF fail at the stage where they produce estrogen (and are pre-pubertal in appearance). **E)** A model showing the possible arrest point where defects in spermatogenesis occur in non-breeding NMRs (indicated by the color graduation in the horizontal bar). Breeding males have the ability to complete all stages of spermatogenesis; however, non-breeding animals fail at postmeiotic stages and have further deficiencies in sperm maturation.

## Additional file legends

**Additional file 1:** Animals/samples used and details of their subsequent analysis.

Tabs in the table indicate:

**A) Sacrificed animals (NMRs) for the experiments**: Table headers indicate animal/sample number in colony, social group (colony) where the animal belonged, year of birth, age (in years), body weight (in gram), whether used for sequencing (RNA-seq) or Immunofluorescene (IF), and the name used in the manuscript.

***B)* RNA-seq information**. Table headers indicate: Tissue, name used to identify sample in manuscript (Name in manuscript), RNA-seq sample number, total sequence obtained (number of reads), length of each read in bp (sequence length), together with percentage GC content of total number of reads.

**Additional file 2:** DEGs between brains of Qs and NBFs.

Tabs in the table indicate:

***All DEGs Qs vs NBFs**:* all DEGs between brains of Qs and NBFs.

***DEG up in Q (Qs vs NBFs):*** DEGs with higher expression in Qs brains.

***DEG down in Qs (Qs vs NBFs):*** DEGs with lower expression in Qs brains.

In all tabs the table headers indicate: the naked mole rat gene id from our transcript assembly (NMR_geneID); the log fold change (logFC); counts per million in log scale (logCPM); p-value (PValue); false discovery rate using Benjamini-Hochberg (FDR); annotation of NMR_geneID to mouse Ensembl gene ID (Ensembl_geneID); and annotation of NMR_geneID to mouse gene name (gene_name).

**Additional file 3:** Result of enrichment analysis on DEGs between brains of Qs and NBFs using Metascape.

The GroupID, ID for clustering of related terms, the enriched terms (Term), the description of the enriched terms (Description), the p-value in log10 scale (p-values calculated based on accumulative hypergeometric distribution), q value in log10 scale (q-values are calculated using the Banjamini-Hochberg procedure to account for multiple testing), list of Symbols of upload hits in this term (Symbols).

**Additional file 4:** Result of enrichment analysis on DEGs between brains of Qs and NBFs using Enrichr (GO Biological Process).

Enrichment analysis was performed as described in material and methods. The table headers indicate: Term; enriched GO Biological Process; p-value, computed by Fisher’s exact test; adjusted p-value, an adjusted p-value using the Benjamini-Hochberg method for correction for multiple hypotheses testing; the rank score or (z-score), computed using a modification to Fisher’s exact test in which a z-score was computed for deviation from an expected rank; combined score, a combination of the p-value and z-score calculated by multiplying the two scores (c = log(p) * z); Genes, belonging to the enriched category.

**Additional file 5:** Result of enrichment analysis on DEGs between brains of Qs and NBFs using Enrichr (GO molecular functions).

The table headers indicate: Term; enriched GO molecular functions; p-value, computed by fisher’s exact test; adjusted p-value, an adjusted p-value using the Benjamini-Hochberg method for correction for multiple hypotheses testing; the rank score or (z-score), computed using a modification to Fisher’s exact test in which a z-score was computed for deviation from an expected rank; combined score, a combination of the p-value and z-score calculated by multiplying the two scores (c = log(p) * z); Genes, belonging to the enriched category.

**Additional file 6:** DEGs between brains of BMs and NBMs.

Tabs in the table indicate:

**All_DEGs (BMs vs NBMs) tab:** all DEGs between brains of BMs vs NBMs.

**DEG_up_in_BMs (BMs vs NBMs) tab**: DEGs with higher expression in BMs brains.

**DEG_down_in_BMs (BMs vs NBMs) tab**: DEGs with lower expression in BMs brains.

In all cases the table headers indicate: the naked mole rat gene id from our transcript assembly (NMR_geneID); the log fold change (logFC); counts per million in log scale (logCPM); pvalue (PValue); false discovery rate using Benjamini-Hochberg (FDR); annotation of NMR_geneID to mouse Ensembl gene ID (Ensembl_geneID); and annotation of NMR_geneID to mouse gene name (gene_name).

**Additional file 7:** Result of enrichment analysis on DEGs between brains of BMs and NBMs using Metascape.

The table headers indicate: the GroupID, ID for clustering of related terms, the enriched terms (Term), the description of the enriched terms (Description), the p-value in log10 scale (p-values calculated based on accumulative hypergeometric distribution), q-value in log10 scale (q-values are calculated using the Banjamini-Hochberg procedure to account for multiple testing), list of Symbols of upload hits in this term (Symbols).

**Additional file 8:** Result of enrichment analysis on DEGs between brains of BMs and NBMs using Enrichr (GO Biological Process).

Enrichment analysis was performed as described in material and methods. The table headers indicate: Term; enriched GO Biological Process; p-value, computed by fisher’s exact test; adjusted P-value, an adjusted p-value using the Benjamini-Hochberg method for correction for multiple hypotheses testing; the rank score or (z-score), computed using a modification to Fisher’s exact test in which a z-score was computed for deviation from an expected rank; combined score, a combination of the p-value and z-score calculated by multiplying the two scores (c = log(p) * z); Genes, belonging to the enriched category.

**Additional file 9:** Result of enrichment analysis on DEGs between brains of BMs and NBMs using Enrichr (GO molecular functions).

The table headers indicate: Term; enriched GO molecular functions; p-value, computed by fisher’s exact test; adjusted p-value, an adjusted p-value using the Benjamini-Hochberg method for correction for multiple hypotheses testing; the rank score or (z-score), computed using a modification to Fisher’s exact test in which a z-score was computed for deviation from an expected rank; combined score, a combination of the p-value and z-score calculated by multiplying the two scores (c = log(p) * z); Genes, belonging to the enriched category.

**Additional file 10:** DEGs between brains of Qs and BMs.

Tabs in the table indicate:

**All_DEGs (Qs_vs_BMs) tab:** all DEGs between brains of Qs and BMs.

**DEG_and_up_in_Q (Qs_vs_BMs) tab:** DEGs with higher expression in Qs brain.

**DEG_and_down_in_Q (Qs_vs_BMs) tab:** DEGs with lower expression in Qs brain.

In all cases, the table headers indicate: the naked mole rat gene id from our transcript assembly (NMR_geneID); the log fold change (logFC); counts per million in log scale (logCPM); p-value (PValue); false discovery rate using Benjamini-Hochberg (FDR); annotation of NMR_geneID to mouse Ensembl gene ID (Ensembl_geneID); and annotation of NMR_geneID to mouse gene name (gene_name).

**Additional file 11:** Result of enrichment analysis on DEGs between brains of Qs and BMs using Metascape.

The table headers indicate: the GroupID, ID for clustering of related terms, the enriched terms (Term), the description of the enriched terms (Description), the p-value in log10 scale (p-values calculated based on accumulative hypergeometric distribution), q-value in log10 scale (q-values are calculated using the Banjamini-Hochberg procedure to account for multiple testing), list of Symbols of upload hits in this term (Symbols).

**Additional file 12:** Result of enrichment analysis on DEGs between brains of Qs and BMs using Enrichr (GO Biological Process).

The table headers indicate: Term; enriched GO Biological Process; p-value, computed by fisher’s exact test; adjusted p-value, an adjusted p-value using the Benjamini-Hochberg method for correction for multiple hypotheses testing; the rank score or (z-score), computed using a modification to Fisher’s exact test in which a z-score was computed for deviation from an expected rank; combined score, a combination of the p-value and z-score calculated by multiplying the two scores (c = log(p) * z); Genes, belonging to the enriched category.

**Additional file 13:** Result of enrichment analysis on DEGs between brains of Qs and BMs using Enrichr (GO molecular functions).

The table headers indicate: Term; enriched GO molecular functions; p-value, computed by fisher’s exact test; adjusted p-value, an adjusted p-value using the Benjamini-Hochberg method for correction for multiple hypotheses testing; the rank score or (z-score), computed using a modification to Fisher’s exact test in which a z-score was computed for deviation from an expected rank; combined score, a combination of the p-value and z-score calculated by multiplying the two scores (c = log(p) * z); Genes, belonging to the enriched category.

**Additional file 14:** DEGs between brains of NBFs and NBMs.

Tabs in the table indicate:

**All_DEGs (NBFs vs NBMs) tab**: all DEGs brains of NBFs and NBMs.

**DEGs up in NBF (NBFs vs NBMs) tab**: DEGs with higher expression in NBFs brain.

**DEGs down in NBF (NBFs vs NBMs) tab:** DEGs with lower expression in NBFs brain.

In all tabs the table headers indicate: the naked mole rat gene id from our transcript assembly (NMR_geneID); the log fold change (logFC); counts per million in log scale (logCPM); p-value (PValue); false discovery rate using Benjamini-Hochberg (FDR); annotation of NMR_geneID to mouse Ensembl gene ID (Ensembl_geneID); and annotation of NMR_geneID to mouse gene name (gene_name).

**Additional file 15:** DEGs between brains of Qs and NBMs.

Tabs in the table indicate:

**All_DEGs (Qs_vs_NBMs) tab:** all DEGs between brains of Qs and NBMs.

**DEG_up_in_Q (Qs_vs_NBMs) tab:** DEGs with higher expression in Qs brain.

**DEG_down_in_Q (Qs_vs_NBMs) tab:** DEGs with lower expression in Qs brain.

In all cases the table headers indicate: the naked mole rat gene id from our transcript assembly (NMR_geneID); the log fold change (logFC); counts per million in log scale (logCPM); p-value (PValue); false discovery rate using Benjamini-Hochberg (FDR); annotation of NMR_geneID to mouse Ensembl gene ID (Ensembl_geneID); and annotation of NMR_geneID to mouse gene name (gene_name).

**Additional file 16**: Defining queen specific genes (Q genes). The list of DEGs in the comparison between Qs vs NBFs, Qs vs NBMs, Qs vs BMs were mapped to each other to identify queen specific genes.

The different tables in the excel sheet are:

**Q vs NBF NBM BM all tab:** DEGs between Qs vs NBFs, Qs vs NBMs and Qs vs BMs.

***Q vs NBF NBM all tab***: all DEGs between Qs vs NBFs and Qs vs NBMs.

***Q vs NBF NBM BM down in Q tab***: genes that are differentially expressed in the comparison (Qs vs NBFs, Qs vs NBMs, and Qs vs BMs) and also show lower expression in Q (down regulation in Q)

***Q vs NBF NBM down in Q tab:*** genes that are differentially expressed in the comparison (Qs vs NBFs, Qs vs NBMs) and also show lower expression in Q (down regulation in Q)

***Q vs NBF NBM BM up in Q tab:*** genes that are differentially expressed in the comparison (Qs vs NBFs, Qs vs NBMs, and Qs vs BMs) and also show higher expression in Q (up regulation in Q)

***Q vs NBF NBM up in Q tab:*** genes that are differentially expressed in the comparison (Qs vs NBFs, Qs vs NBMs) and also show higher expression in Q (up regulation in Q)

**Additional file 17:** Result of enrichment analysis of Q genes using Metascape.

The table headers indicate: GroupID, ID for clustering of related terms, the enriched terms (Term), the description of the enriched terms (Description), the p-value in log10 scale (p-values calculated based on accumulative hypergeometric distribution), q value in log10 scale (q-values are calculated using the Banjamini-Hochberg procedure to account for multiple testing), list of Symbols of upload hits in this term (Symbols).

**Additional file 18:** Result of enrichment analysis of Q genes using Enrichr (GO Biological Process).

The table headers indicate: the Term; enriched GO Biological Process; P-value, computed by fisher’s exact test; adjusted p-value, an adjusted p-value using the Benjamini-Hochberg method for correction for multiple hypotheses testing; the rank score or (z-score), computed using a modification to Fisher’s exact test in which a z-score was computed for deviation from an expected rank; combined score, a combination of the p-value and z-score calculated by multiplying the two scores (c = log(p) * z); Genes, belonging to the enriched category.

**Additional file 19:** Result of enrichment analysis of Q genes using Enrichr (GO molecular functions).

The table headers indicate: the Term; enriched GO molecular functions; p-value, computed by fisher’s exact test; adjusted P-value, an adjusted p-value using the Benjamini-Hochberg method for correction for multiple hypotheses testing; the rank score or (z-score), computed using a modification to Fisher’s exact test in which a z-score was computed for deviation from an expected rank; combined score, a combination of the p-value and z-score calculated by multiplying the two scores (c = log(p) * z); Genes, belonging to the enriched category.

**Additional file 20:** DEGs between brains of BMs and NBFs.

Tabs in the table indicate:

**All DEGs BMs vs NBFs tab:** all DEGs brains of BMs and NBFs.

**DEGs Up in BM (BMs vs NBFs) tab:** DEGs with higher expression in BMs brain.

**DEGs down in BM (BMs vs NBFs) tab:** DEGs with lower expression in BMs brain.

In all cases the table headers indicate: the naked mole rat gene id from our transcript assembly (NMR_geneID); the log fold change (logFC); counts per million in log scale (logCPM); p-value (PValue); false discovery rate using Benjamini-Hochberg (FDR); annotation of NMR_geneID to mouse Ensembl gene ID (Ensembl_geneID); and annotation of NMR_geneID to mouse gene name (gene_name).

**Additional file 21:** Defining BM specific genes (BM genes). The list of DEGs in the comparison between BMs vs NBFs, BMs vs NBMs, were mapped to each other to identify breeding male specific genes.

**Additional file 22:** Result of enrichment analysis of BM genes using Metascape.

Enrichment analysis was performed as described in material and methods. The output of the enrichment analysis is presented here. The table headers indicate: The GroupID, ID for clustering of related terms, the enriched terms (Term), the description of the enriched terms (Description), the p-value in log10 scale (p-values calculated based on accumulative hypergeometric distribution), q value in log10 scale (q-values are calculated using the Banjamini-Hochberg procedure to account for multiple testing), list of Symbols of upload hits in this term (Symbols).

**Additional file 23:** Result of enrichment analysis of BM genes using Enrichr (GO Biological Process).

The table headers indicate: the Term; enriched GO Biological Process; p-value, computed by fisher’s exact test; adjusted p-value, an adjusted p-value using the Benjamini-Hochberg method for correction for multiple hypotheses testing; the rank score or (z-score), computed using a modification to Fisher’s exact test in which a z-score was computed for deviation from an expected rank; combined score, a combination of the p-value and z-score calculated by multiplying the two scores (c = log(p) * z); Genes, belonging to the enriched category.

**Additional file 24:** Result of enrichment analysis of BM genes using Enrichr (GO molecular functions).

The table headers indicate: the Term; enriched GO molecular functions; p-value, computed by fisher’s exact test; adjusted p-value, an adjusted p-value using the Benjamini-Hochberg method for correction for multiple hypotheses testing; the rank score or (z-score), computed using a modification to Fisher’s exact test in which a z-score was computed for deviation from an expected rank; combined score, a combination of the p-value and z-score calculated by multiplying the two scores (c = log(p) * z); Genes, belonging to the enriched category.

**Additional file 25:** DEGs between Qs and NBFs ovary.

Tabs in the table indicate:

***All_DEGs (Qs_vs_NBFs) tab:*** all DEGs between ovaries of Qs and NBFs.

***DEG_and_down_in_Q (Qs_vs_NBFs) tab:*** DEGs with lower expression in Qs ovary.

***DEG_and_up_in_Q (Qs_vs_NBFs) tab:*** DEGs with lower expression in in Qs ovary.

In all tabs the table headers indicate: the naked mole rat gene id from our transcript assembly (NMR_geneID); the log fold change (logFC); counts per million in log scale (logCPM); p-value (PValue); false discovery rate using Benjamini-Hochberg (FDR); annotation of NMR_geneID to mouse Ensembl gene ID (Ensembl_geneID); and annotation of NMR_geneID to mouse gene name (gene_name).

**Additional file 26:** Result of enrichment analysis on DEGs between Qs and NBFs ovary using Metascape. The table headers indicate: The GroupID, ID for clustering of related terms, the enriched terms (Term), the description of the enriched terms (Description), the p-value in log10 scale (p-values calculated based on accumulative hypergeometric distribution), q value in log10 scale (q-values are calculated using the Banjamini-Hochberg procedure to account for multiple testing), list of symbols of upload hits in this term (Symbols).

The tabs in the table show:

**A) all DEGs:** enriched pathways, Biological Process, and molecular functions for all DEGs between Qs and NBFs ovary.

**B) DEGs up in Q:** enriched pathways, Biological Process, and molecular functions for DEGs between Qs and NBFs ovary, and up regulated in Q compared to NBF.

**C) DEGs down in Q:** enriched pathways, Biological Process, and molecular functions for DEGs between Qs and NBFs ovary, and down regulated in Q compared to NBF.

**Additional file 27:** Result of enrichment analysis on DEGs between Qs and NBFs ovary using Enrichr (GO Biological Process).

The table headers indicate: the Term; enriched GO Biological Process; p-value, computed by fisher’s exact test; adjusted p-value, an adjusted p-value using the Benjamini-Hochberg method for correction for multiple hypotheses testing; the rank score or (z-score), computed using a modification to Fisher’s exact test in which a z-score was computed for deviation from an expected rank; combined score, a combination of the p-value and z-score calculated by multiplying the two scores (c = log(p) * z); Genes, belonging to the enriched category.

The tabs in the table show:

**A) All DEGs:** enriched Biological Process for all DEGs between Qs and NBFs ovary.

**B) DEGs down in Q:** enriched Biological Process for DEGs between Q and NBFs ovary, and down regulated in Qs compared to NBFs.

**C) DEGs up in Q:** enriched Biological Process for DEGs between Qs and NBFs ovary, and up regulated in Qs compared to NBFs.

**Additional file 28:** Result of enrichment analysis on DEGs between Qs and NBFs ovary using Enrichr (GO molecular functions).

The table headers indicate: the Term; enriched GO molecular functions; p-value, computed by fisher’s exact test; adjusted p-value, an adjusted p-value using the Benjamini-Hochberg method for correction for multiple hypotheses testing; the rank score or (z-score), computed using a modification to Fisher’s exact test in which a z-score was computed for deviation from an expected rank; combined score, a combination of the p-value and z-score calculated by multiplying the two scores (c = log(p) * z); Genes, belonging to the enriched category.

The tabs in the table show:

**A) All DEGs:** enriched molecular functions for all DEGs between Qs and NBFs ovary.

**B) DEGs down in Qs:** enriched molecular functions for DEGs between Qs and NBFs ovary, and down regulated in Qs compared to NBFs.

**C) DEGs up in Qs:** enriched Biological Process for DEGs between Qs and NBFs ovary, and up regulated in Qs compared to NBFs.

**Additional file 29:** DEGs between BMs and NBMs testis.

Tabs in the table indicate:

***All DEGs* (BMs vs NBMs) *tab*:** all DEGs between BMs and NBMs testis.

***DEG down in BM* (BMs vs NBMs) *tab*:** DEGs and higher expression in BMs testis.

***DEG up in BM* (BMs vs NBMs) *tab*:** DEGs and lower expression in BMs testis.

**Germonline genes:** Gene list obtained form Germonline (germonline.org)

In all tabs the table headers indicate: the naked mole rat gene id from our transcript assembly (NMR_geneID); the log fold change (logFC); counts per million in log scale (logCPM); pvalue (PValue); false discovery rate using Benjamini-Hochberg (FDR); annotation of NMR_geneID to mouse Ensembl gene ID (Ensembl_geneID); and annotation of NMR_geneID to mouse gene name (gene_name).

**Additional file 30:** Result of enrichment analysis on DEGs between BMs and NBMs testis using Metascape.

The table headers indicate: the GroupID, ID for clustering of related terms, the enriched terms (Term), the description of the enriched terms (Description), the p-value in log10 scale (p-values calculated based on accumulative hypergeometric distribution), q-value in log10 scale (q-values are calculated using the Banjamini-Hochberg procedure to account for multiple testing), list of Symbols of upload hits in this term (Symbols).

The tabs in the table show:

**A) All DEGs:** enriched pathways, Biological Process, and molecular functions for all DEGs between BMs and NBM testis.

**B) DEGs down in_BM:** enriched pathways, Biological Process, and molecular functions for DEGs between BMs and NBMs testis, and up regulated in BMs compared to NBMs.

**C) DEGs up in BM:** enriched pathways, Biological Process, and molecular functions for DEGs between BMs and NBMs testis, and up regulated in BMs compared to NBMs.

**D) reprod &steroid respon:** selected genes that belong to terms, multicellular organism reproduction, response to steroid hormone, multicellular organism reproduction and response to steroid hormone.

**Additional file 31:** Result of enrichment analysis on differentially expressed between BMs and NBMs testis using Enrichr (GO Biological Process).

The table headers indicate: Term; enriched GO Biological Process; p-value, computed by fisher’s exact test; adjusted p-value, an adjusted p-value using the Benjamini-Hochberg method for correction for multiple hypotheses testing; the rank score or (z-score), computed using a modification to Fisher’s exact test in which a z-score was computed for deviation from an expected rank; combined score, a combination of the p-value and z-score calculated by multiplying the two scores (c = log(p) * z); Genes, belonging to the enriched category.

The tabs in the table show:

**A) All_deg:** GO Biological Process enrichment for all DEGs between BMs and NBMs testis.

**B) DEGs up in BM:** GO Biological Process enrichment for DEGs between BMs and NBMs testis, and up regulated in BMs compared to NBMs.

**C) DEGs down in BM:** GO Biological Process enrichment for DEGs between BMs and NBMs testis, and down regulated in BMs compared to NBMs.

**Additional file 32:** Result of enrichment analysis on DEGs between BMs and NBMs testis using Enrichr (GO molecular functions). The table headers indicate: Term; enriched GO molecular functions; p-value, p-value, computed by fisher’s exact test; adjusted p-value, an adjusted p-value using the Benjamini-Hochberg method for correction for multiple hypotheses testing; the rank score or (z-score), computed using a modification to Fisher’s exact test in which a z-score was computed for deviation from an expected rank; combined score, a combination of the p-value and z-score calculated by multiplying the two scores (c = log(p) * z); Genes, belonging to the enriched category.

The tabs in the table show:

**A) All DEG:** GO molecular functions enrichment for DEGs between BMs and NBMs testis.

**B) DEG down in BM:** GO molecular functions enrichment for DEGs between BMs and NBMs testis, and down regulated in BMs compared to NBMs.

**C) DEG up in BM:** GO molecular functions enrichment for DEGs between BMs and NBMs testis, and up regulated in BMs compared to NBMs.

## References

[1] Szathmary E, Smith JM: The major evolutionary transitions. Nature 1995, 374:227–232.

[2] Nowak MA, Tarnita CE, Wilson EO: The evolution of eusociality. Nature 2010, 466:1057–1062.

[3] Wilson EO: The insect societies. Cambridge, Mass.: Belknap Press of Harvard University Press; 1971.

[4] Michener CD: Comparative Social Behavior of Bees. Annual Review of Entomology 1969, 14:299–342

[5] Crespi BJ, Yanega D: The definition of eusociality. Oxford Journals, Behavioral Ecology 1995, 6:109–115.

[6] Duffy JE, Morrison CL, Rios R: Multiple origins of eusociality among sponge-dwelling shrimps (Synalpheus). Evolution 2000, 54:503–516.

[7] Duffy JE, Macdonald KS: Kin structure, ecology and the evolution of social organization in shrimp: a comparative analysis. Proc Biol Sci 2010, 277:575–584.

[8] Jarvis JU: Eusociality in a mammal: cooperative breeding in naked mole-rat colonies. Science 1981, 212:571–573.

[9] Friedman DA, Gordon DM: Ant Genetics: Reproductive Physiology, Worker Morphology, and Behavior. Annu Rev Neurosci 2016, 39:41–56.

[10] J FE, H K: On Biomass and Trophic Structure of the Central Amazonian Rain Forest Ecosystem Biotropica 1973, 5:2–14.

[11] Hölldobler B, Wilson EO: The ants. Cambridge, Mass.: Belknap Press of Harvard University Press; 1990.

[12] M OR, CG F: African mole-rats: eusociality, relatedness and ecological constraints. In Ecology of social evolution. Edited by J H, J K. Berlin, Germany: Springer; 2008: 205–220

[13] Faulkes CG, Bennett NC, Bruford MW, O’Brien HP, Aguilar GH, Jarvis JU: Ecological constraints drive social evolution in the African mole-rats. Proc Biol Sci 1997, 264:1619–1627.

[14] M. JJU, C. BN: Eusociality has evolved independently in two genera of bathyergid mole-rats — but occurs in no other subterranean mammal. Behavioral Ecology and Sociobiology 1993, 33:253–260.

[15] RA. B: The ecology of naked mole-rat colonies: burrowing, food, and limiting factors. In The Biology of the Naked Mole-Rat. Edited by PW S, JUM J, RD A. New Jersey: Princeton University Press; 1991: 97–136

[16] EA. L, PW. S: Social organization of naked mole-rat colonies: Evidence for divisions of labor. In The Biology of the Naked Mole-Rat. Edited by PW S, JUM J, RD A. New Jersey: Princeton University Press; 1991: 275–336

[17] Buffenstein R: The naked mole-rat: a new long-living model for human aging research. J Gerontol A Biol Sci Med Sci 2005, 60:1369–1377.

[18] Faulkes CG, Abbott DH, Jarvis JU: Social suppression of ovarian cyclicity in captive and wild colonies of naked mole-rats, Heterocephalus glaber. J Reprod Fertil 1990, 88:559–568.

[19] Faulkes CG, Abbott DH, Jarvis JU, Sherriff FE: LH responses of female naked mole-rats, Heterocephalus glaber, to single and multiple doses of exogenous GnRH. J Reprod Fertil 1990, 89:317–323.

[20] Faulkes CG, Abbott DH: Social control of reproduction in breeding and non-breeding male naked mole-rats (Heterocephalus glaber). J Reprod Fertil 1991, 93:427–435.

[21] Faulkes CG, Abbott DH: Evidence that primer pheromones do not cause social suppression of reproduction in male and female naked mole-rats (Heterocephalus glaber). J Reprod Fertil 1993, 99:225–230.

[22] Clarke FM, Faulkes CG: Dominance and queen succession in captive colonies of the eusocial naked mole-rat, Heterocephalus glaber. Proc Biol Sci 1997, 264:993–1000.

[23] Clarke FM, Faulkes CG: Hormonal and behavioural correlates of male dominance and reproductive status in captive colonies of the naked mole-rat, Heterocephalus glaber. Proc Biol Sci 1998, 265:1391–1399.

[24] Margulis SW, Saltzman W, Abbott DH: Behavioral and hormonal changes in female naked mole-rats (Heterocephalus glaber) following removal of the breeding female from a colony. Horm Behav 1995, 29:227–247.

[25] O’Riain MJ, Jarvis JU, Alexander R, Buffenstein R, Peeters C: Morphological castes in a vertebrate. Proc Natl Acad Sci U S A 2000, 97:13194–13197.

[26] Dengler-Crish CM, Catania KC: Phenotypic plasticity in female naked mole-rats after removal from reproductive suppression. J Exp Biol 2007, 210:4351–4358.

[27] Faulkes CG, Abbott DH, Jarvis JU: Social suppression of reproduction in male naked mole-rats, Heterocephalus glaber. J Reprod Fertil 1991, 91:593–604.

[28] Faulkes CG, Trowell SN, Jarvis JU, Bennett NC: Investigation of numbers and motility of spermatozoa in reproductively active and socially suppressed males of two eusocial African mole-rats, the naked mole-rat (Heterocephalus glaber) and the Damaraland mole-rat (Cryptomys damarensis). J Reprod Fertil 1994, 100:411–416.

[29] Legan SJ, Karsch FJ: Neuroendocrine regulation of the estrous cycle and seasonal breeding in the ewe. Biol Reprod 1979, 20:74–85.

[30] Glasier A, McNeilly AS, Baird DT: Induction of ovarian activity by pulsatile infusion of LHRH in women with lactational amenorrhoea. Clin Endocrinol (Oxf) 1986, 24:243–252.

[31] Abbott DH, Hodges JK, George LM: Social status controls LH secretion and ovulation in female marmoset monkeys (Callithrix jacchus). J Endocrinol 1988, 117:329–339.

[32] Goodman RL, Lehman MN: Kisspeptin neurons from mice to men: similarities and differences. Endocrinology 2012, 153:5105–5118.

[33] Zhou S, Holmes MM, Forger NG, Goldman BD, Lovern MB, Caraty A, Kallo I, Faulkes CG, Coen CW: Socially regulated reproductive development: analysis of GnRH-1 and kisspeptin neuronal systems in cooperatively breeding naked mole-rats (Heterocephalus glaber). J Comp Neurol 2013, 521:3003–3029.

[34] Peragine DE, Pokarowski M, Mendoza-Viveros L, Swift-Gallant A, Cheng HM, Bentley GE, Holmes MM: RFamide-related peptide-3 (RFRP-3) suppresses sexual maturation in a eusocial mammal. Proc Natl Acad Sci U S A 2017, 114:1207–1212.

[35] Toth AL, Varala K, Henshaw MT, Rodriguez-Zas SL, Hudson ME, Robinson GE: Brain transcriptomic analysis in paper wasps identifies genes associated with behaviour across social insect lineages. Proc Biol Sci 2010, 277:2139–2148.

[36] Ferreira PG, Patalano S, Chauhan R, Ffrench-Constant R, Gabaldon T, Guigo R, Sumner S: Transcriptome analyses of primitively eusocial wasps reveal novel insights into the evolution of sociality and the origin of alternative phenotypes. Genome Biol 2013, 14:R20.

[37] Standage DS, Berens AJ, Glastad KM, Severin AJ, Brendel VP, Toth AL: Genome, transcriptome and methylome sequencing of a primitively eusocial wasp reveal a greatly reduced DNA methylation system in a social insect. Mol Ecol 2016, 25:1769–1784.

[38] Simola DF, Wissler L, Donahue G, Waterhouse RM, Helmkampf M, Roux J, Nygaard S, Glastad KM, Hagen DE, Viljakainen L, et al. Social insect genomes exhibit dramatic evolution in gene composition and regulation while preserving regulatory features linked to sociality. Genome Res 2013, 23:1235–1247.

[39] Kapheim KM, Pan H, Li C, Salzberg SL, Puiu D, Magoc T, Robertson HM, Hudson ME, Venkat A, Fischman BJ, et al. Social evolution. Genomic signatures of evolutionary transitions from solitary to group living. Science 2015, 348:1139–1143.

[40] Sadd BM, Barribeau SM, Bloch G, de Graaf DC, Dearden P, Elsik CG, Gadau J, Grimmelikhuijzen CJ, Hasselmann M, Lozier JD, et al. The genomes of two key bumblebee species with primitive eusocial organization. Genome Biol 2015, 16:76.

[41] Smith CR, Toth AL, Suarez AV, Robinson GE: Genetic and genomic analyses of the division of labour in insect societies. Nat Rev Genet 2008, 9:735–748.

[42] Hunt GJ, Gadau JR: Editorial: Advances in Genomics and Epigenomics of Social Insects. Front Genet 2016, 7:199.

[43] Yu C, Li Y, Holmes A, Szafranski K, Faulkes CG, Coen CW, Buffenstein R, Platzer M, de Magalhaes JP, Church GM: RNA sequencing reveals differential expression of mitochondrial and oxidation reduction genes in the long-lived naked mole-rat when compared to mice. PLoS One 2011, 6:e26729.

[44] Davies KT, Bennett NC, Tsagkogeorga G, Rossiter SJ, Faulkes CG: Family Wide Molecular Adaptations to Underground Life in African Mole-Rats Revealed by Phylogenomic Analysis. Mol Biol Evol 2015, 32:3089–3107.

[45] Fang X, Seim I, Huang Z, Gerashchenko MV, Xiong Z, Turanov AA, Zhu Y, Lobanov AV, Fan D, Yim SH, et al. Adaptations to a subterranean environment and longevity revealed by the analysis of mole rat genomes. Cell Rep 2014, 8:1354–1364.

[46] Holmes MM, Goldman BD, Goldman SL, Seney ML, Forger NG: Neuroendocrinology and sexual differentiation in eusocial mammals. Front Neuroendocrinol 2009, 30:519–533.

[47] Bjorklund A, Dunnett SB: Dopamine neuron systems in the brain: an update. Trends Neurosci 2007, 30:194–202.

[48] Ben-Jonathan N, Hnasko R: Dopamine as a prolactin (PRL) inhibitor. Endocr Rev 2001, 22:724–763.

[49] Jung D, Kee K: Insights into female germ cell biology: from in vivo development to in vitro derivations. Asian J Androl 2015, 17:415–420.

[50] Bukovsky A, Caudle MR, Svetlikova M, Wimalasena J, Ayala ME, Dominguez R: Oogenesis in adult mammals, including humans: a review. Endocrine 2005, 26:301–316.

[51] Reiss M, Sants H: Behaviour and Social Organisation. 1987.

[52] Pan H, O’Brien M J, Wigglesworth K, Eppig JJ, Schultz RM: Transcript profiling during mouse oocyte development and the effect of gonadotropin priming and development in vitro. Dev Biol 2005, 286:493–506.

[53] Pan L, Gong W, Zhou Y, Li X, Yu J, Hu S: A comprehensive transcriptomic analysis of infant and adult mouse ovary. Genomics Proteomics Bioinformatics 2014, 12:239–248.

[54] Simpson ER, Zhao Y, Agarwal VR, Michael MD, Bulun SE, Hinshelwood MM, Graham-Lorence S, Sun T, Fisher CR, Qin K, Mendelson CR: Aromatase expression in health and disease. Recent Prog Horm Res 1997, 52:185–213; discussion 213-184.

[55] Simpson ER, Mahendroo MS, Means GD, Kilgore MW, Hinshelwood MM, Graham-Lorence S, Amarneh B, Ito Y, Fisher CR, Michael MD, et al.: Aromatase cytochrome P450, the enzyme responsible for estrogen biosynthesis. Endocr Rev 1994, 15:342–355.

[56] Griswold MD: Spermatogenesis: The Commitment to Meiosis. Physiol Rev 2016, 96:1–17.

[57] Griswold MD, Oatley JM: Concise review: Defining characteristics of mammalian spermatogenic stem cells. Stem Cells 2013, 31:8–11.

[58] J W, JB S, CM L: Mammalian spermatogenesis. Func Dev Embryol 2007, 1:99–117.

[59] Ramaswamy S, Weinbauer GF: Endocrine control of spermatogenesis: Role of FSH and LH/ testosterone. Spermatogenesis 2014, 4: e996025.

[60] Themmen APN, Huhtaniemi IT: Mutations of gonadotropins and gonadotropin receptors: elucidating the physiology and pathophysiology of pituitary-gonadal function. Endocr Rev 2000, 21:551–583.

[61] Gospodarowicz D: Properties of the luteinizing hormone receptor of isolated bovine corpus luteum plasma membranes. J Biol Chem 1973, 248:5042–5049.

[62] Rivero-Muller A, Potorac I, Pintiaux A, Daly AF, Thiry A, Rydlewski C, Nisolle M, Parent AS, Huhtaniemi I, Beckers A: A novel inactivating mutation of the LH/chorionic gonadotrophin receptor with impaired membrane trafficking leading to Leydig cell hypoplasia type 1. Eur J Endocrinol 2015, 172:K27–36.

[63] Ascoli M, Fanelli F, Segaloff DL: The lutropin/choriogonadotropin receptor, a 2002) perspective. Endocr Rev 2002, 23:141–174.

[64] Sassone-Corsi P: Unique chromatin remodeling and transcriptional regulation in spermatogenesis. Science 2002, 296:2176–2178.

[65] Steger K: Transcriptional and translational regulation of gene expression in haploid spermatids. Anat Embryol (Berl) 1999, 199:471–487.

[66] Balhorn R: The protamine family of sperm nuclear proteins. Genome Biol 2007, 8:227.

[67] Steger K, Failing K, Klonisch T, Behre HM, Manning M, Weidner W, Hertle L, Bergmann M, Kliesch S: Round spermatids from infertile men exhibit decreased protamine-1 and −2 mRNA. Hum Reprod 2001, 16:709–716.

[68] Bjorndahl L, Kvist U: Human sperm chromatin stabilization: a proposed model including zinc bridges. Mol Hum Reprod 2010, 16:23–29.

[69] Carrell DT, Emery BR, Hammoud S: Altered protamine expression and diminished spermatogenesis: what is the link? Hum Reprod Update 2007, 13:313–327.

[70] Luke L, Vicens A, Tourmente M, Roldan ER: Evolution of protamine genes and changes in sperm head phenotype in rodents. Biol Reprod 2014, 90:67.

[71] Imken L, Rouba H, El Houate B, Louanjli N, Barakat A, Chafik A, McElreavey K: Mutations in the protamine locus: association with spermatogenic failure? Mol Hum Reprod 2009, 15:733–738.

[72] Ravel C, Chantot-Bastaraud S, El Houate B, Berthaut I, Verstraete L, De Larouziere V, Lourenco D, Dumaine A, Antoine JM, Mandelbaum J, et al. Mutations in the protamine 1 gene associated with male infertility. Mol Hum Reprod 2007, 13:461–464.

[73] Clark AG, Civetta A: Evolutionary biology. Protamine wars. Nature 2000, 403:261, 263.

[74] Petersen C, Aumuller G, Bahrami M, Hoyer-Fender S: Molecular cloning of Odf3 encoding a novel coiled-coil protein of sperm tail outer dense fibers. Mol Reprod Dev 2002, 61:102–112.

[75] Baltz JM, Williams PO, Cone RA: Dense fibers protect mammalian sperm against damage. Biol Reprod 1990, 43:485–491.

[76] Miki K, Willis WD, Brown PR, Goulding EH, Fulcher KD, Eddy EM: Targeted disruption of the Akap4 gene causes defects in sperm flagellum and motility. Dev Biol 2002, 248:331–342.

[77] Moretti E, Scapigliati G, Pascarelli NA, Baccetti B, Collodel G: Localization of AKAP4 and tubulin proteins in sperm with reduced motility. Asian J Androl 2007, 9:641–649.

[78] Lardenois A, Gattiker A, Collin O, Chalmel F, Primig M: GermOnline 4.0 is a genomics gateway for germline development, meiosis and the mitotic cell cycle. Database (Oxford) 2010, 2010:baq030.

[79] Gasbarri A, Sulli A, Packard MG: The dopaminergic mesencephalic projections to the hippocampal formation in the rat. Prog Neuropsychopharmacol Biol Psychiatry 1997, 21:1–22.

[80] Kempadoo KA, Mosharov EV, Choi SJ, Sulzer D, Kandel ER: Dopamine release from the locus coeruleus to the dorsal hippocampus promotes spatial learning and memory. Proc Natl Acad Sci U S A 2016, 113:14835–14840.

[81] Waddell S: Dopamine reveals neural circuit mechanisms of fly memory. Trends Neurosci 2010, 33:457–464.

[82] Wicker-Thomas C, Hamann M: Interaction of dopamine, female pheromones, locomotion and sex behavior in Drosophila melanogaster. J Insect Physiol 2008, 54:1423–1431.

[83] Cottrell GA: Occurrence of dopamine and noradrenaline in the nervous tissue of some invertebrate species. Br J Pharmacol Chemother 1967, 29:63–69.

[84] Kindt KS, Quast KB, Giles AC, De S, Hendrey D, Nicastro I, Rankin CH, Schafer WR: Dopamine mediates context-dependent modulation of sensory plasticity in C. elegans. Neuron 2007, 55:662–676.

[85] Carlsson A: Thirty years of dopamine research. Adv Neurol 1993, 60:1–10.

[86] Schultz W: Responses of midbrain dopamine neurons to behavioral trigger stimuli in the monkey. J Neurophysiol 1986, 56:1439–1461.

[87] Joshua M, Adler A, Bergman H: The dynamics of dopamine in control of motor behavior. Curr Opin Neurobiol 2009, 19:615–620.

[88] Dayan P, Balleine BW: Reward, motivation, and reinforcement learning. Neuron 2002, 36:285–298.

[89] Wise RA: Dopamine, learning and motivation. Nat Rev Neurosci 2004, 5:483–494.

[90] Heinz A, Schlagenhauf F: Dopaminergic dysfunction in schizophrenia: salience attribution revisited. Schizophr Bull 2010, 36:472–485.

[91] Sasaki K, Yamasaki K, Nagao T: Neuro-endocrine correlates of ovarian development and egg-laying behaviors in the primitively eusocial wasp (Polistes chinensis). J Insect Physiol 2007, 53:940–949.

[92] Penick CA, Brent CS, Dolezal K, Liebig J: Neurohormonal changes associated with ritualized combat and the formation of a reproductive hierarchy in the ant Harpegnathos saltator. J Exp Biol 2014, 217:1496–1503.

[93] Okada Y, Sasaki K, Miyazaki S, Shimoji H, Tsuji K, Miura T: Social dominance and reproductive differentiation mediated by dopaminergic signaling in a queenless ant. J Exp Biol 2015, 218:1091–1098.

[94] Bloch G, Simon T, Robinson GE, Hefetz A: Brain biogenic amines and reproductive dominance in bumble bees (Bombus terrestris). J Comp Physiol A 2000, 186:261–268.

[95] JW. H, Woodring J: Elevated brain dopamine levels associated with ovary development in queenless worker honeybees (Apis mellifera L) Comparative Biochemistry and Physiology 1995, 111C:271–279.

[96] T. D, Simoes ZLP, Bitondi MMG: Dietary dopamine causes ovary activation in queenless Apis mellifera workers. Apidologie, Springer Verlag 2003, 34:281–289.

[97] Matsuyama S, Nagao T, Sasaki K: Consumption of tyrosine in royal jelly increases brain levels of dopamine and tyramine and promotes transition from normal to reproductive workers in queenless honey bee colonies. Gen Comp Endocrinol 2015, 211:1–8.

[98] Beggs KT, Glendining KA, Marechal NM, Vergoz V, Nakamura I, Slessor KN, Mercer AR: Queen pheromone modulates brain dopamine function in worker honey bees. Proc Natl Acad Sci U S A 2007, 104:2460–2464.

[99] Akasaka S, Sasaki K, Harano K, Nagao T: Dopamine enhances locomotor activity for mating in male honeybees (Apis mellifera L.). J Insect Physiol 2010, 56:1160–1166.

[100] Mezawa R, Akasaka S, Nagao T, Sasaki K: Neuroendocrine mechanisms underlying regulation of mating flight behaviors in male honey bees (Apis mellifera L.). Gen Comp Endocrinol 2013, 186:108–115.

[101] S. DOT, A. R, W. E: Reactivation of juvenile hormone synthesis in adult drones of the honey bee, Apis mellifera carnica. Experientia, Springer 1995, 51:945–952.

[102] Giray T, Robinson GE: Common endocrine and genetic mechanisms of behavioral development in male and worker honey bees and the evolution of division of labor. Proc Natl Acad Sci U S A 1996, 93:11718–11722.

[103] Fitzgerald P, Dinan TG: Prolactin and dopamine: what is the connection? A review article. J Psychopharmacol 2008, 22:12–19.

[104] Brown RS, Herbison AE, Grattan DR: Effects of Prolactin and Lactation on A15 Dopamine Neurones in the Rostral Preoptic Area of Female Mice. J Neuroendocrinol 2015, 27:708–717.

[105] Kauppila A, Martikainen H, Puistola U, Reinila M, L R: Hypoprolactinemia and ovarian function. Fertility and Sterility 1988, 49:437–441.

[106] Majumdar A, Mangal NS: Hyperprolactinemia. J Hum Reprod Sci 2013, 6:168–175.

[107] Brown RS, Herbison AE, Grattan DR: Prolactin regulation of kisspeptin neurones in the mouse brain and its role in the lactation-induced suppression of kisspeptin expression. J Neuroendocrinol 2014, 26:898–908.

[108] Snowdon CT, Ziegler TE: Variation in prolactin is related to variation in sexual behavior and contact affiliation. PLoS One 2015, 10:e0120650.

[109] Fisher CR, Graves KH, Parlow AF, Simpson ER: Characterization of mice deficient in aromatase (ArKO) because of targeted disruption of the cyp19 gene. Proc Natl Acad Sci U S A 1998, 95:6965–6970.

[110] Toda K, Takeda K, Okada T, Akira S, Saibara T, Kaname T, Yamamura K, Onishi S, Shizuta Y: Targeted disruption of the aromatase P450 gene (Cyp19) in mice and their ovarian and uterine responses to 17betaoestradiol. J Endocrinol 2001, 170:99–111.

[111] Hewitt SC, Winuthayanon W, Korach KS: What’s new in estrogen receptor action in the female reproductive tract. J Mol Endocrinol 2016, 56:R55–71.

[112] R. L: What is an hystricomorph? In The Biology of hystricomorph rodents. Edited by IW R, Weir BJ e. London: Zoological Society of London; 1974: 7–20

[113] Kent J, Ryle M: Histochemical studies on three gonadotrophin-responsive enzymes in the infantile mouse ovary. J Reprod Fertil 1975, 42:519–536.

[114] Paranko J, Pelliniemi LJ: Differentiation of smooth muscle cells in the fetal rat testis and ovary: localization of alkaline phosphatase, smooth muscle myosin, F-actin, and desmin. Cell Tissue Res 1992, 268:521–530.

[115] Brannstrom M, Mayrhofer G, Robertson SA: Localization of leukocyte subsets in the rat ovary during the periovulatory period. Biol Reprod 1993, 48:277–286.

[116] Best CL, Pudney J, Welch WR, Burger N, Hill JA: Localization and characterization of white blood cell populations within the human ovary throughout the menstrual cycle and menopause. Hum Reprod 1996, 11:790–797.

[117] Magoffin DA: The ovarian androgen-producing cells: a 2001) perspective. Rev Endocr Metab Disord 2002, 3:47–53.

[118] Berkholtz CB, Lai BE, Woodruff TK, Shea LD: Distribution of extracellular matrix proteins type I collagen, type IV collagen, fibronectin, and laminin in mouse folliculogenesis. Histochem Cell Biol 2006, 126:583–592.

[119] S. A: FastQC: a quality control tool for high throughput sequence data. Babraham Bioinformatics 2010.

[120] Bolger AM, Lohse M, Usadel B: Trimmomatic: a flexible trimmer for Illumina sequence data. Bioinformatics 2014, 30:2114–2120.

[121] Kim D, Pertea G, Trapnell C, Pimentel H, Kelley R, Salzberg SL: TopHat2: accurate alignment of transcriptomes in the presence of insertions, deletions and gene fusions. Genome Biol 2013, 14:R36.

[122] Langmead B, Salzberg SL: Fast gapped-read alignment with Bowtie 2. Nat Methods 2012, 9:357–359.

[123] Trapnell C, Williams BA, Pertea G, Mortazavi A, Kwan G, van Baren MJ, Salzberg SL, Wold BJ, Pachter L: Transcript assembly and quantification by RNA-Seq reveals unannotated transcripts and isoform switching during cell differentiation. Nat Biotechnol 2010, 28:511–515.

[124] Yates A, Akanni W, Amode MR, Barrell D, Billis K, Carvalho-Silva D, Cummins C, Clapham P, Fitzgerald S, Gil L, et al. Ensembl 2016. Nucleic Acids Res 2016, 44:D710–716.

[125] Camacho C, Coulouris G, Avagyan V, Ma N, Papadopoulos J, Bealer K, Madden TL: BLAST+: architecture and applications. BMC Bioinformatics 2009, 10:421.

[126] Altschul SF, Gish W, Miller W, Myers EW, Lipman DJ: Basic local alignment search tool. J Mol Biol 1990, 215:403–410.

[127] Anders S, Pyl PT, Huber W: HTSeq--a Python framework to work with high-throughput sequencing data. Bioinformatics 2015, 31:166–169.

[128] Risso D, Schwartz K, Sherlock G, Dudoit S: GC-content normalization for RNA-Seq data. BMC Bioinformatics 2011, 12:480.

[129] Risso D, Ngai J, Speed TP, Dudoit S: Normalization of RNA-seq data using factor analysis of control genes or samples. Nat Biotechnol 2014, 32:896–902.

[130] Robinson MD, McCarthy DJ, Smyth GK: edgeR: a Bioconductor package for differential expression analysis of digital gene expression data. Bioinformatics 2010, 26:139–140.

[131] Tripathi S, Pohl MO, Zhou Y, Rodriguez-Frandsen A, Wang G, Stein DA, Moulton HM, DeJesus P, Che J, Mulder LC, et al. Meta- and Orthogonal Integration of Influenza “OMICs” Data Defines a Role for UBR4 in Virus Budding. Cell Host Microbe 2015, 18:723–735.

[132] Kuleshov MV, Jones MR, Rouillard AD, Fernandez NF, Duan Q, Wang Z, Koplev S, Jenkins SL, Jagodnik KM, Lachmann A, et al. Enrichr: a comprehensive gene set enrichment analysis web server 2016) update. Nucleic Acids Res 2016, 44:W90–97.

